# Single cell transcriptional and chromatin accessibility profiling redefine cellular heterogeneity in the adult human kidney

**DOI:** 10.1101/2020.06.14.151167

**Authors:** Yoshiharu Muto, Parker C. Wilson, Haojia Wu, Sushrut S. Waikar, Benjamin D. Humphreys

## Abstract

The integration of single cell transcriptome and chromatin accessibility datasets enables a deeper understanding of cell heterogeneity. We performed single nucleus ATAC (snATAC-seq) and RNA (snRNA-seq) sequencing to generate paired, cell-type-specific chromatin accessibility and transcriptional profiles of the adult human kidney. We demonstrate that snATAC-seq is comparable to snRNA-seq in the assignment of cell identity and can further refine our understanding of functional heterogeneity in the nephron. The majority of differentially accessible chromatin regions are localized to promoters and a significant proportion are closely-associated with differentially expressed genes. Cell-type-specific enrichment of transcription factor binding motifs implicates the activation of NFκB that promotes *VCAM1* expression and drives transition between a subpopulation of proximal tubule epithelial cells. These datasets can be visualized at this resource: http://humphreyslab.com/SingleCell/. Our multi-omics approach improves the ability to detect unique cell states within the kidney and redefines cellular heterogeneity in the proximal tubule and thick ascending limb.

## Introduction

Single cell RNA sequencing (scRNA-seq) has fostered a greater understanding of the genes and pathways that define cell identity in the kidney^1^. Multiple scRNA-seq atlases of mature human^2–5^ and mouse kidney ^6,7^ have established how transcription contributes to cell type specificity. Recent methods have expanded this approach to single cell profiling of chromatin accessibility ^8–10^. Single cell assay for transposase-accessible chromatin using sequencing (scATAC-seq) is an extension of bulk ATAC-seq^11^ that employs hyperactive Tn5 transposase to measure chromatin accessibility in thousands of individual cells^9^. Joint profiling by scRNA-seq and scATAC-seq in the adult mouse kidney has provided a framework for understanding how chromatin accessibility regulates transcription^8^, however, the single-cell epigenomic landscape of the human kidney has not been described.

Integration and analysis of multimodal single cell datasets is an emerging field with enormous potential for accelerating our understanding of kidney disease and development^12,13^. Bioinformatics tools can extract unique information from scATAC-seq datasets that is otherwise unavailable by scRNA-seq. Prediction of cell-type-specific cis-regulatory DNA interactions^14^ and transcription factor activity^15^ are two methods that complement the transcriptional information obtained by scRNA-seq. Long-range chromatin-chromatin interactions play an important role in transcriptional regulation and are influenced by transcription factor binding^16,17^. Chromatin accessibility profiling will help to identify distant regulatory regions that influence transcription via long-range interactions.

The kidney is composed of diverse cell types with distinct subpopulations and single cell sequencing can dissect cellular heterogeneity at high resolution^1^. For example, a small subset of cells within the proximal tubule and Bowman’s capsule express vimentin, CD24, and CD133^18,19^. These cells have a distinct morphology and expression profile and undergo expansion after acute kidney injury^19^. Furthermore, they have reported potential for tubular and podocyte regeneration ^20–23^ and have been implicated in the development of renal cell carcinoma ^24^. However, the sparsity of these cells has hampered further characterization. Another example of cellular heterogeneity is seen in the thick ascending limb (TAL). Studies in mouse and human suggest that there are structural and functional differences between medullary and cortical TAL, however, the signaling pathways that drive these differences are not well defined ^25^.

Chromatin accessibility is a dynamic process that drives nephron development^26^. Nephron progenitors have distinct chromatin accessibility profiles that change as they differentiate^10,26^. The role of chromatin accessibility in promotion or inhibition of kidney repair and regeneration has important implications for designing therapies for acute and chronic kidney disease^27^ and may help to improve directed differentiation of kidney organoids^28^.

We have performed single nucleus ATAC (snATAC-seq) and RNA (snRNA-seq) sequencing to examine how chromatin accessibility can refine our understanding of cell state and function in the mature human kidney. We generated an interactive multimodal atlas encompassing both transcriptomic and epigenomic data (http://humphreyslab.com/SingleCell/). Combined snRNA-seq and snATAC-seq analysis improved our ability to detect unique cell states within the proximal tubule and thick ascending limb and redefines cellular heterogeneity that may contribute to kidney regeneration and segment-specific cation permeability.

## Results

### Single cell transcriptional and chromatin accessibility profiling in the adult human kidney

snRNA-seq and snATAC-seq was performed on 5 healthy adult kidney samples (Fig. 1a). Selected patients ranged in age from 50 to 62 years and included men (N=3) and women (N=2). All patients had preserved kidney function (mean sCr = 1.07 mg/dl, eGFR = 64.4 +/− 4.7 ml/min/1.73m^2^). Histologic review showed no significant glomerulosclerosis or interstitial fibrosis and tubular atrophy (Supplementary Table 1). We performed snRNA-seq to determine the cellular composition of our samples, annotate cells based on their transcriptional profiles, and inform our snATAC-seq analysis (Fig. 1a). snRNA-seq identified all major cell types within the kidney cortex (Fig. 1b, Supplementary Fig. 1a) based on expression of lineage-specific markers (Fig. 1c, Supplementary Fig. 1b) ^29^. We detected proximal tubule (PT), parietal epithelial cells (PEC), loop of Henle (TAL), distal tubule (DCT1, DCT2), connecting tubule (CNT), collecting duct (PC, ICA, ICB), endothelial cells (ENDO), glomerular cell types (MES, PODO), fibroblasts (FIB), and a small population of leukocytes (LEUK) (Supplementary Table 2, Supplementary Data 1). Notably, there was a subpopulation of proximal tubule that had increased expression of *VCAM1* (PT_VCAM1). This subpopulation also expressed *HAVCR1* (kidney injury molecule-1), which is a biomarker for acute kidney injury and predictor of long-term renal outcomes^30,31^.

**Figure 1.**
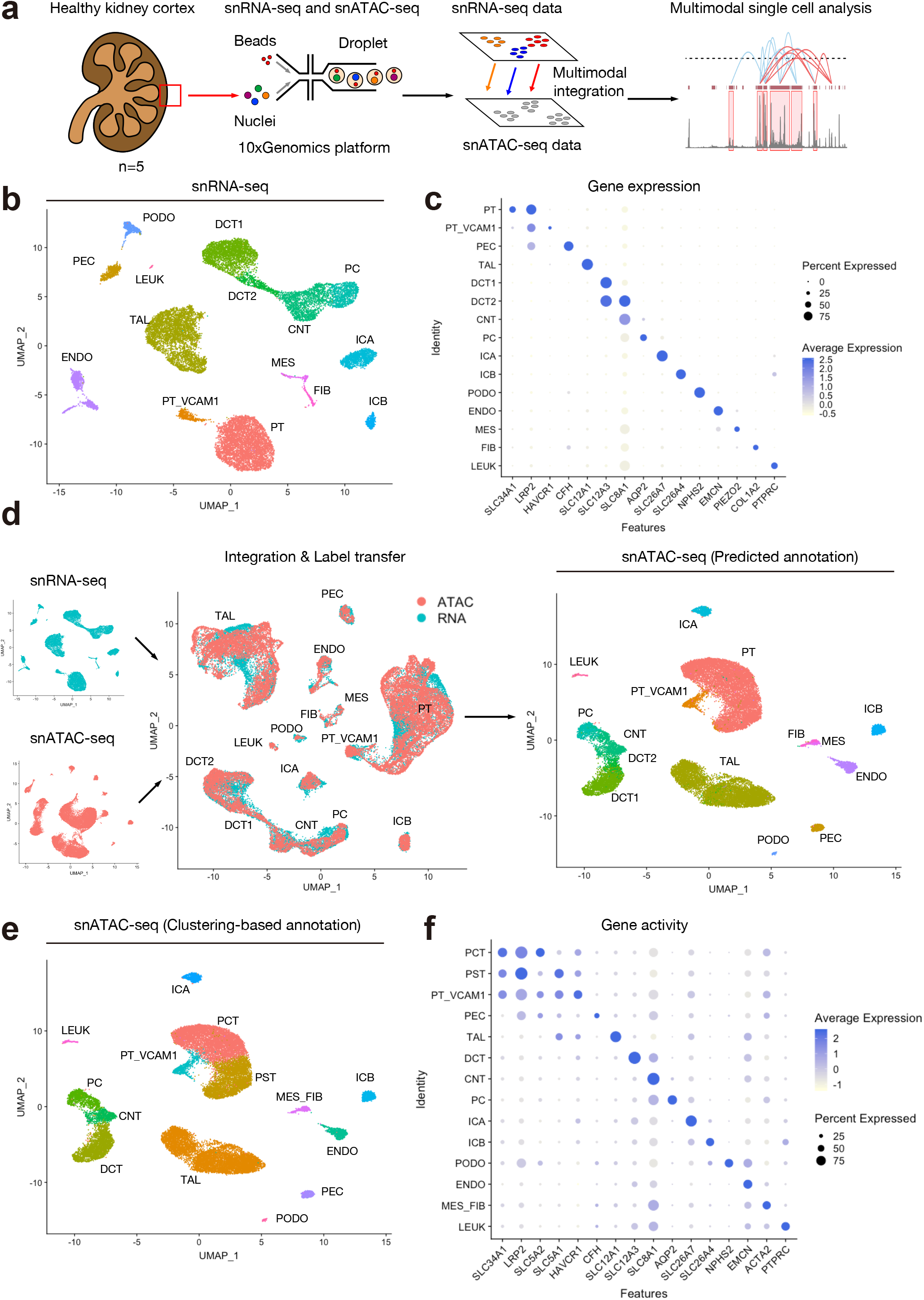
Single Cell Transcriptional and Chromatin Accessibility Profiling on the human adult kidneys. (**a**) Graphical abstract of experimental methodology. n=5 human adult kidneys were analyzed with snRNA-seq and snATAC-seq. (**b**) Umap plots of snRNA-seq dataset. PT, proximal tubule; PT_VCAM1, subpopulation of proximal tubule with VCAM1 expression; PEC, parietal epithelial cells; TAL, thick ascending limb of loop of Henle; DCT, distal convoluted tubule; CNT, connecting tubule; PC, principle cells, ICA, Type A intercalated cells; ICB, Type B intercalated cells; PODO, podocyte; ENDO, endothelial cells; MES, mesangial cells, FIB, fibroblasts; LEUK, leukocytes. (**c**) Dotplots of snRNA-seq dataset showing gene expression patterns of cluster-enriched markers. (**d**) Multi-omics integration strategy for processing the snATAC-seq dataset. Following integration and label transfer, the snATAC-seq dataset was filtered using a 97% prediction score threshold for cell type assignment. (**e**) Umap plots of snATAC-seq dataset with gene activities-based cell type assignments. PCT, proximal convoluted tubule; PST, proximal straight tubule. (**f**) Dotplots of snATAC-seq dataset showing gene activity patterns of cell type markers.

### Integration of single nucleus RNA and ATAC datasets for prediction and validation of ATAC cell type assignments

snATAC-seq captures the chromatin accessibility profile of individual cells^9^. Relatively less is known about cell-type-specific chromatin accessibility profiles; so we leveraged our annotated snRNA-seq dataset to predict snATAC-seq cell types with label transfer^12^. Label transfer was performed by creating a gene activity matrix from the snATAC-seq data, which is a measure of chromatin accessibility within the gene body and promoter of protein-coding genes. Transfer anchors were identified between the ‘reference’ snRNA-seq dataset and ‘query’ gene activity matrix followed by assignment of predicted cell types. The distribution of snATAC-seq prediction scores showed that the vast majority of cells had a high prediction score and were confidently assigned to a single cell type (Supplementary Fig. 2). The snATAC-seq dataset was filtered using a 97% confidence threshold for cell type assignment to remove heterotypic doublets. Comparison between snATAC-seq cell type predictions obtained by label transfer (Fig. 1d) and curated annotations of unsupervised clusters (Fig. 1e, f, Supplementary Fig. 1c, d and Supplementary Table 3) indicates that all major cell types were present in both datasets and that snATAC-seq is comparable to snRNA-seq in the detection and assignment of cell identities (Supplementary Fig. 3). We performed downstream analyses with gene-activity-based cell type assignments, which were obtained by unsupervised clustering of the snATAC-seq dataset. Interestingly, snATAC-seq was able to detect two subpopulations within the proximal tubule cluster, which likely represent the proximal convoluted tubule (Fig. 1e, PCT) and the proximal straight tubule (Fig. 1e, PST). PCT showed greater chromatin accessibility in *SLC5A2,* which encodes sodium glucose cotransporter 2 (SGLT2); whereas PST showed greater accessibility in *SLC5A1* (Fig. 1f, Supplementary Fig. 4). SGLT2 reabsorbs glucose in the S1 and S2 segments of the proximal tubule and SGLT1 (*SLC5A1*) is located in S3^32^. The delineation between S1/S2 and S3 was less clear in the snRNA-seq dataset (Supplementary Fig. 4), which suggests that snATAC-seq provides complementary information that may refine cell type assignment; particularly for genes transcribed at low levels or genes that are not detected by snRNA-seq. The chromatin accessibility profile of S1/S2 may be of clinical interest in determining the factors that drive glucose reabsorption, which is the therapeutic target of SGLT2 inhibitors^32^. Together, our multimodal snATAC-seq and snRNA-seq analysis improved our ability to dissect cellular heterogeneity.

### Chromatin accessibility defines cell type

We detected 214,890 accessible chromatin regions among 27,034 cells in the snATAC-seq library. We compared these regions to a previously-published dataset of DNase I-hypersensitive sites (DHS) in bulk glomeruli and tubulointerstitium^33^. DNase hypersensitivity is an alternative measure of chromatin accessibility and approximately 50% of all regions identified by our pipeline were overlapping with a DHS in the glomerulus or tubulointerstitium. The proportion of overlapping regions increased to ~85% when our dataset was filtered for regions contained in at least 10% of nuclei (Supplementary Fig. 5). These data suggest that snATAC-seq is a robust method for the detection of accessible chromatin in the adult kidney.

We used the R package Signac^12^ to investigate differences in chromatin accessibility between cell types. Cell types can be distinguished based on whether differentially accessible chromatin regions (DAR) are ‘open’ or ‘closed’ (Fig. 2a, Supplementary Data 2). Approximately 20% (mean proportion=0.203 +/− 0.04) of DAR were closely-associated with differentially expressed genes in their respective cell types (Supplementary Table 4). For example, *LRP2* is a lineage-specific gene expressed in the proximal tubule and a coverage plot in this region shows an increase in number and amplitude of ATAC peaks within its promoter and gene body (Fig. 2a). In fact, the majority of DAR were located in a promoter region within 3kb of the nearest transcriptional start site (Fig. 2b, Supplementary Fig. 6). The second most common location was intronic and the distribution of DAR was relatively conserved across cell types (Fig. 2c). A minority of cell-type-specific differentially expressed genes were closely-associated with a DAR (mean proportion=0.358 +/− 0.07), which raises the question of assigning function to DAR that are not located near differentially expressed genes. Regulatory regions can associate via long-range interactions and a DAR does not necessarily regulate the closest gene^34^. Bioinformatics tools can infer regulatory chromatin interactions and may be useful for assigning function to DAR^14^. Long-range interactions mediate the association between enhancers and promoters via chromatin looping and are regulated in part by transcription factors^16^.

**Figure 2.**
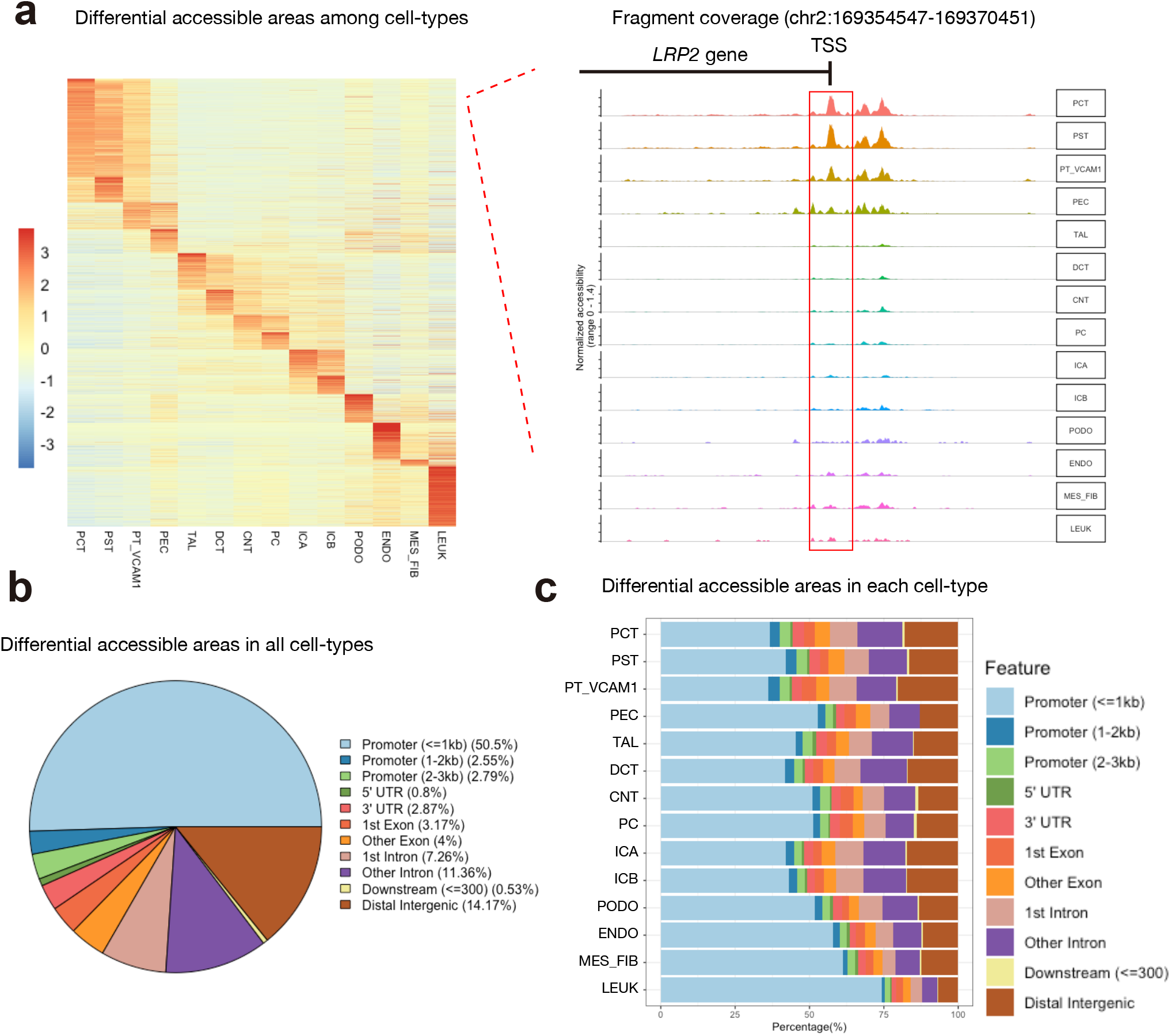
Distribution of cell type-specific chromatin accessible regions. (**a**) Heatmap of average number of Tn5 cut sites within a differentially accessible region (DAR) for each cell type (left). Fragment coverage (frequency of Tn5 insertion) around the DAR (DAR +/−50 Kb) on the LRP2 gene promoter is shown (right). (**b**) Pie plot of genomic annotations for all DAR in the dataset. (C) Bar plot of annotated DAR locations for each cell type.

### Chromatin accessibility is associated with cell-type-specific transcription factor activity and chromatin interaction networks

Transcription factors are key determinants of cell fate that drive cellular differentiation in kidney ageing and development^35–37^. Transcription factor ‘activity’ can be predicted for individual cell types based on the presence of binding motifs within differentially accessible chromatin regions (DAR). We used chromVAR^15^ to infer transcription-factor-associated chromatin accessibility in our snATAC-seq dataset. We observed that individual cell types can be defined by transcription factor ‘activity’ (Fig. 3a, Supplementary Data 3), suggesting that cell-type-specific transcription factors likely regulate chromatin accessibility. For example, *HNF4A* encodes a key transcription factor that drives proximal tubule differentiation^38^. chromVAR detected an enrichment of HNF4A binding motifs within DAR in the proximal tubule (Fig. 3b, motif activity) that was supported by increased chromatin accessibility in *HNF4A* (Fig. 3b, gene activity) and increased *HNF4A* transcription in the snRNA-seq dataset (Fig. 3b, gene expression). A similar pattern was seen for *TFAP2B,* which regulates development in the distal nephron^39^. There was increased TFAP2B transcription factor ‘activity’ in the thick ascending limb and distal convoluted tubule (Fig. 3b, motif activity), in addition to increased chromatin accessibility in *TFAP2B* (Fig. 3b, gene activity) and increased *TFAP2B* transcription (Fig. 3b, gene expression).

**Figure 3.**
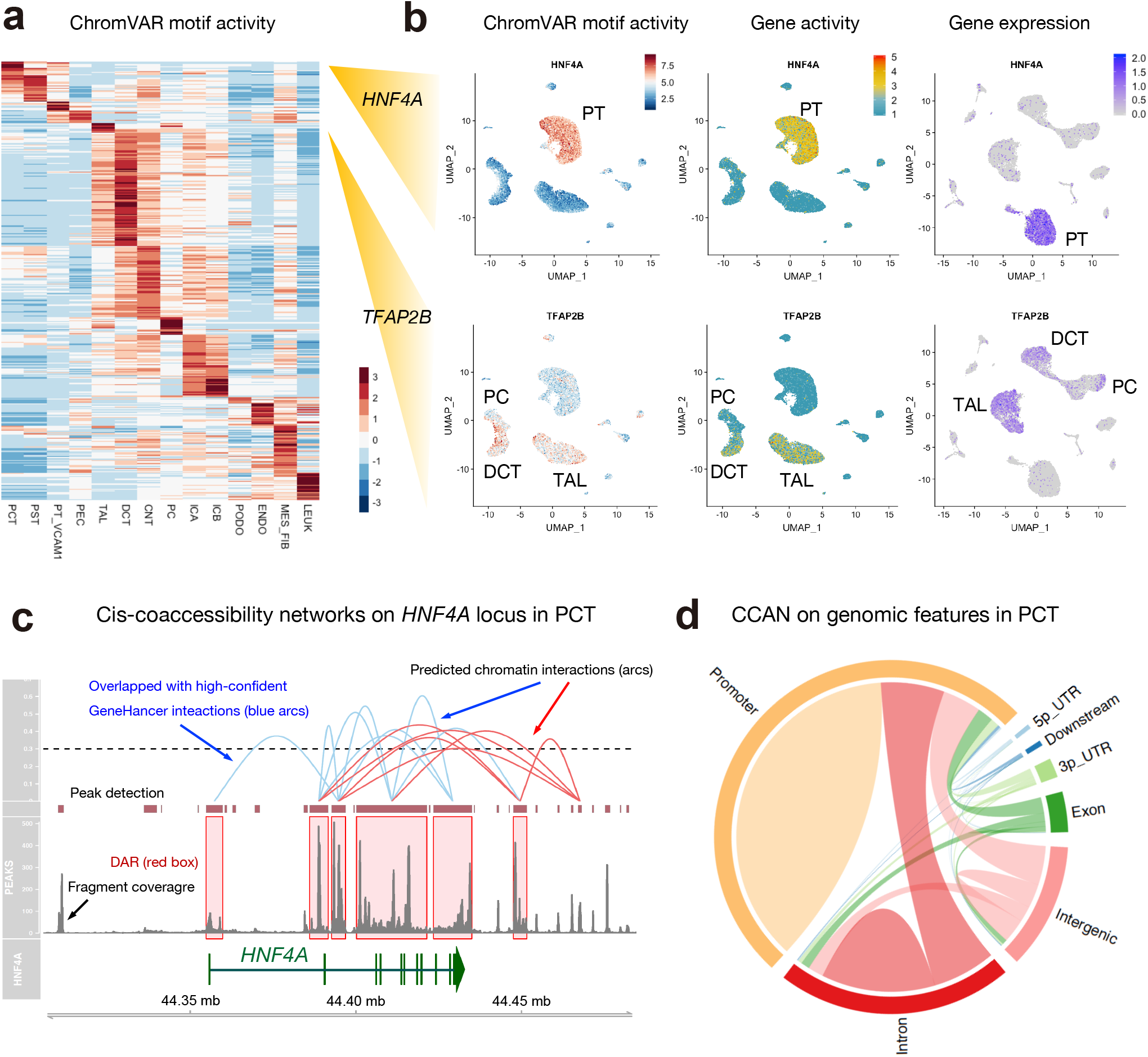
Cell-type-specific transcription factor activity and chromatin interaction networks. (**a**) Heatmap of average chromVar motif activity for each cell type. (**b**) Umap plot displaying chromVAR motif activity (left), gene activity (middle) and gene expression (right) of *HNF4A* or *TFAP2B*. (**c**) Cis-coaccessibility networks (CCAN, red or blue arcs) near the *HNF4A* locus in the proximal convoluted tubule with multiple connections between differentially accessible regions (red boxes). DAR overlapping with high-confidence GeneHancer interactions are shown as blue arcs. Fragment coverage (frequency of Tn5 insertion) and called ATAC peaks are shown in the lower half. *HNF4A* gene track is shown along the bottom of the image. (**d**) Genomic features of Cis-coaccessibility networks (CCAN) in the PCT.

We used the R package Cicero^14^ to predict cis-regulatory chromatin interactions for individual cell types. Cis-coaccessibility networks (CCAN) are families of chromatin-chromatin interactions that regulate gene expression by approximating enhancers and promoters. Within *HNF4A,* we observed a robust CCAN in the proximal convoluted tubule with multiple connections (red or blue arcs) between differentially accessible regions (Fig. 3c, red boxes) in the promoter, gene body, and distal 3’ region (Fig. 3c). We compared these interactions with the GeneHancer database^40^ to determine which connections had been previously-reported in the literature. GeneHancer is a collection of human enhancers and their inferred target genes created using 4 methods: promoter capture Hi-C, enhancer-targeted transcription factors, expression quantitative trait loci, and tissue co-expression correlation between genes and enhancer RNA. The subset of GeneHancer interactions with ‘double elite’ status is the most stringent set of interactions in the database^40^ and the majority of predicted Cicero interactions within 50kb of a cell-type-specific differentially accessible region (DAR) were overlapping with ‘double elite’ GeneHancer interactions (Fig. 3c, blue arcs). The proportion of Cicero connections present in the ‘double elite’ GeneHancer database was dependent on the Cicero coaccessibility score, which is a measure of increased confidence of the predicted interaction (Supplementary Fig. 7). The Cicero connections with a lower coaccessibility score (threshold=0.1, mean proportion in GeneHancer=0.68) were less likely (p < 0.0001, Paired t-test) to be in the GeneHancer ‘double elite’ database compared to Cicero connections with a higher coaccessibility score (threshold=0.5, mean proportion in GeneHancer=0.75). Within the proximal convoluted tubule, the majority of Cicero connections were either within a promoter region or between a promoter and another location (Fig. 3d) and this distribution was similar in other cell types (Supplementary Fig. 8). In summary, Cicero is a robust method for predicting chromatin-chromatin interactions that may play a role in the regulation of cell-type-specific chromatin accessibility.

### Multi-modal analysis highlights cellular heterogeneity in the thick ascending limb

The thick ascending limb regulates extracellular fluid volume, urinary concentration, and calcium and magnesium homeostasis in the outer medulla and cortex^41^. The majority of bivalent cations are reabsorbed in the cortical segment and are regulated by expression of claudins^42^. Claudins are a family of tight junction proteins that confer segment-specific cation permeability and regulate the reabsorption of Na+, K+, Cl−, Mg++ and Ca++^43^. Claudin-10 expression is enriched in the medullary TAL, whereas claudin-16 is expressed predominantly in the cortical TAL^44^. Pathogenic germline variants in *CLDN16* are causative of familial hypomagnesemia, hypercalciuria and nephrocalcinosis, which can lead to end-stage renal disease^45,46^. Interestingly, *Cldn10* deletion causes hypermagnesemia in mice^47^, and even rescues *Cldn16-* deficient mice from hypomagnesemia and hypercalciuria^48^. These data indicate that CLDN10 and CLDN16 may confer opposing segment-specific tight junction cation selectivity. To determine if we could detect subpopulations of cells with variable claudin expression patterns, we performed unsupervised clustering on the thick ascending limb in our snRNA-seq dataset (Supplementary Fig. 9a) to identify 3 groups of cells. There was a group of cells (*SLC12A1*+*UMOD*+) that expressed cortical TAL markers (*CLDN16*, *KCNJ10* and *PTH1R*) and a second group that expressed the medullary TAL marker, *CLDN10* (Supplementary Fig. 9b). The third group of cells was identified as ascending thin limb (ATL) based on expression of previously-published markers^2^.

We analyzed the thick ascending limb cluster in the snATAC-seq dataset and identified 3 groups of cells that echoed our findings in the snRNA-seq dataset (Supplementary Fig. 9c,d). Our findings suggest that TAL subpopulations can be defined by either transcription or chromatin accessibility profiles. We identified differential transcription factor ‘activity’ for HNF1B and ESRRB that defined cortical and medullary subpopulations (Supplementary Fig. 9e). Furthermore, HNF1B and ESRRB transcription factor motifs were enriched in the differentially accessible chromatin regions (DAR) that distinguish between cortical and medullary thick ascending limb populations (Supplementary Fig. 9f). ESRRB is an orphan nuclear receptor with a critical role in early development and pluripotency^49,50^ and HNF1B is a homeodomain-containing transcription factor that regulates nephrogenesis. Pathogenic germline *HNF1B* variants are a known cause of autosomal dominant tubulointerstitial kidney disease with hypomagnesemia and hypercalciuria^51^. Interestingly, genetic deletion of *Hnf1b* in mouse kidney was found to increase *Cldn10* expression^52^, suggesting that HNF1B may be involved in the regulation of *CLDN10*. Collectively, our multimodal analysis demonstrates heterogeneity within the TAL at a transcriptomic and chromatin accessibility level and highlights transcription factors that likely contribute to these differences.

### NF-kB regulates the molecular signature of a subpopulation of proximal tubule that expresses *VCAM1*

We detected a subset of proximal tubule cells that had increased expression and chromatin accessibility of *VCAM1,* which we designated PT_VCAM1 (Fig. 1). Immunofluorescence studies demonstrated *VCAM1* expression in a scattered distribution amongst proximal tubule epithelium (Fig. 4a). Our single cell studies estimate that PT_VCAM1 represents ~2% of total cells and 6% of proximal tubular epithelium. Despite the fact that kidney samples originated from patients without kidney injury, the PT_VCAM1 population showed increased expression of kidney injury molecule-1 (*HAVCR1,* KIM1), which is a biomarker that is increased in acute kidney injury^30^ and chronic kidney disease^53^. Interestingly, PT_VCAM1 also expressed *VIM* (vimentin), *CD24,* and *CD133* (*PROM1*) (Supplementary Fig. 10), which is consistent with a previously-described population of cells with progenitor-like features in the proximal tubule^18–20^. These findings suggest that the PT_VCAM1 cluster may represent an injured or regenerative subpopulation.

**Figure 4.**
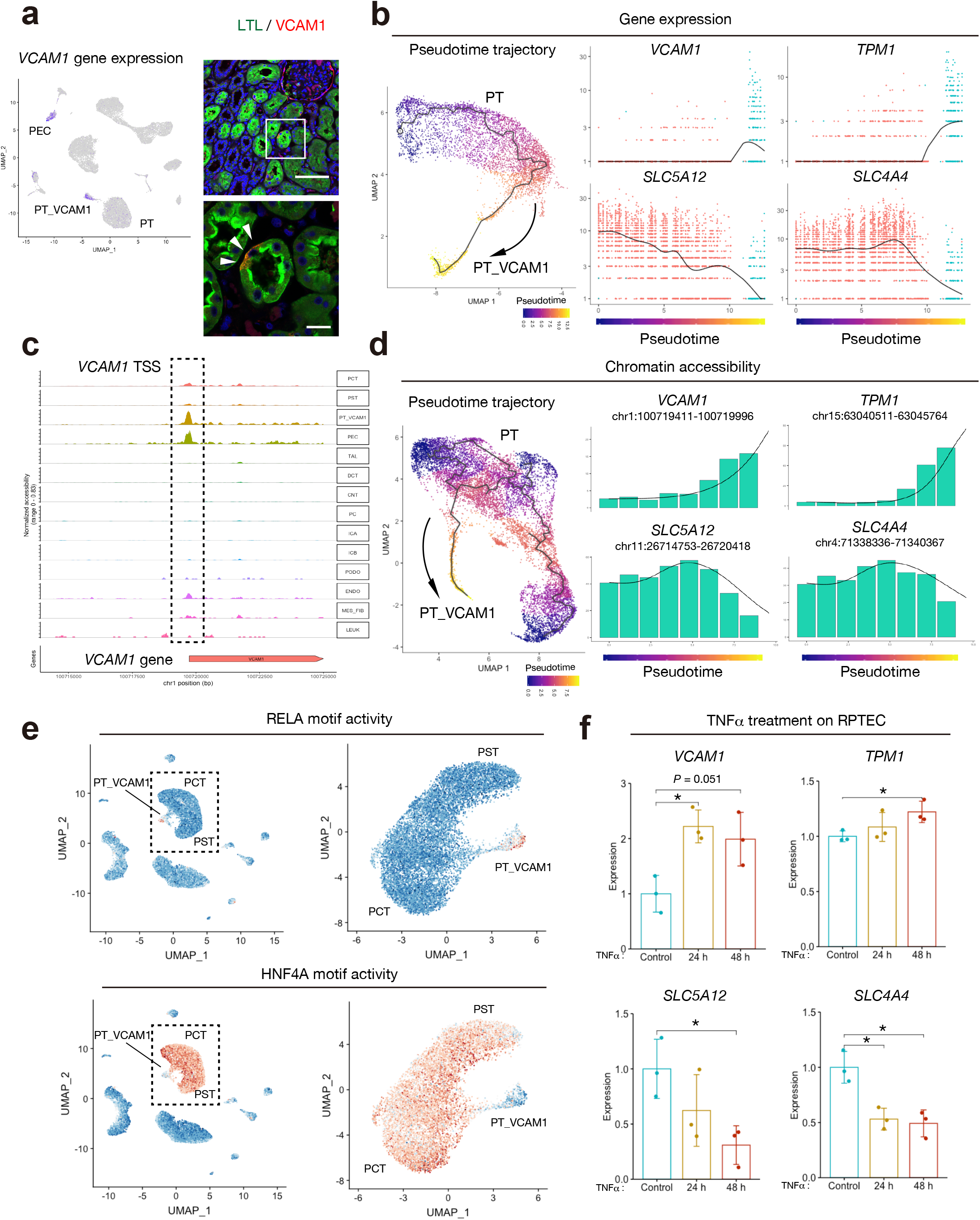
Identification and characterization of previously unrecognized PT subpopulation. (**a**) Umap plot displaying *VCAM1* gene expression in the snRNA-seq dataset (left), and representative immunohistochemical images of VCAM1 (red) or LTL (Lotus tetragonolobus lectin, green) in the adult kidney (n = 3 patients). Arrowheads indicate the VCAM1+ PT. *VCAM1* was expressed in PEC and a subpopulation of LTL+ PT. Scale bar indicates 100 μm (upper right) or 20 μm (lower right). (**b**) Pseudotemporal trajectory from PT to PT_VCAM1 using snRNA-seq was generated with Monocle3 (left), and gene expression dynamics along a pseudotime trajectory from PT to PT_VCAM 1 are shown (right); *VCAM1* (upper left), *TPM1* (upper right), *SLC5A12* (lower left) and *SLC4A4* (lower right). (**c**) Fragment coverage (frequency of Tn5 insertion) around the representative DAR (DAR +/−5000 bp) in *VCAM1* locus. (**d**) Pseudotime trajectory from PT to PT_VCAM1 using snATAC-seq was generated with Cicero (right). Chromatin accessibility dynamics along the pseudotemporal trajectory from PT to PT_VCAM1 are shown (left). chr1:100719411-100719996 (*VCAM1* promoter, upper left); chr15:63040511-63045764 (*TPM1* promoter, upper right), chr11:26714753-26720418 (*SLC5A12* gene body, lower left) and chr4:71338336-71340367 (*SLC4A1* gene body, lower right). (**e**) Feature plot of single cell chromVAR motif activity of RELA and HNF4A in the entire dataset or PT/PT_VCAM1 subset. (f) RT and real-time PCR analysis of mRNAs for *VCAM1*, *TPM1, SLC5A12* and *SLC4A4* in RPTEC treated with TNFα (100 ng/ml) for 24 h or 48 h. \**P* < 0.05 (Student’s *t* test). Bar graphs represent the mean and error bars are the s.d.

We compared the transcriptional profile of PT_VCAM1 to the remaining proximal tubule to identify differentially expressed genes (Supplementary Data 4). Gene ontology enrichment analysis of the differentially expressed genes showed an enrichment for pathways involved in metabolism, cell migration, angiogenesis, proliferation, and apoptosis. In particular, there was enrichment for genes that control branching morphogenesis of epithelial tubes and the MAPK and Wnt signaling pathways (Supplementary Data 5). These results suggest that the signaling pathways in this subpopulation are distinct from the remaining proximal tubule.

We performed pseudotemporal ordering with Monocle^54^ to determine which genes drive the transition from healthy proximal tubule to the PT_VCAM1 state (Fig. 4b). We identified *VCAM1* and *TPM1* as genes that show increased expression in PT_VCAM1 cells and *SLC5A12* and *SLC5A4* as genes that show decreased expression (Fig. 4b). *VCAM1* is a key mediator of angiogenesis^55^ and *TPM1* encodes tropomyosin 1, which is an actin-binding protein involved in the cytoskeletal contraction^56^. In contrast, *SLC5A12* and *SLC4A4* encode a lactate and bicarbonate transporter. SLC5A12 and SLC5A4 are abundantly-expressed in the proximal tubule and VCAM1 and TPM1 can be detected in a subset of cells. We constructed a complementary pseudotemporal trajectory with Cicero (13) to examine changes in chromatin accessibility during the transition from PT to PT_VCAM1. Increased transcription of *VCAM1* and *TPM1* (Fig. 4b) was associated with increased chromatin accessibility within the *VCAM1* gene body and promoter region (Fig. 4c,d). Similarly, decreased transcription of *SLC5A12* and *SLC4A4* (Fig. 4b) was associated with decreased chromatin accessibility (Fig. 4d, Supplementary Fig. 11). We identified transcription factors that likely regulate the transition between proximal tubule and PT_VCAM1 by assessing chromVAR transcription factor activities. Interestingly, the proximal tubule showed robust activity of HNF4A, which was decreased in the PT_VCAM1 cluster and coincided with increased activity of REL and RELA (Fig. 4e). NF-κB is a family of inducible transcription factors that share homology in the Rel domain^57^ that has been implicated in the inflammatory response in renal disease^58^. In particular, ischemia-reperfusion injury-induced acute kidney injury activates NF-κB and NF-κB inhibition improves renal function^59^. Consistent with this finding, gene set enrichment analysis^60,61^ of the differentially expressed genes in PT_VCAM1 compared to PT implicated NF-κB signaling (Supplementary Fig. 12). The TNF family of cytokines are well-established activators of NF-κB ^62^. To further validate these insights, we used TNFα to induce NF-κB in an *in vitro* model of the proximal tubule (RPTEC). TNFα stimulation increased expression of *VCAM1* and *TPM1* and decreased expression of *SLC5A12* and *SLC4A4* (Fig. 4f). These observations suggest that NF-κB induction may play a role in the transition from proximal tubule to the PT_VCAM1 cell state.

### The proportion of PT_VCAM1 is elevated in acute kidney injury and chronic kidney disease

We performed deconvolution of bulk RNA-seq obtained from mouse IRI to determine if the proportion of PT_VCAM1 is related to acute kidney injury. BisqueRNA estimates cell type abundance from bulk RNA-seq using a scRNA-seq reference-based deconvolution ^63^. The estimated proportion of PT_VCAM1 significantly increased 24 hours post-IRI and persisted for at least 7 days; corresponding with a decrease in the proportion of normal PT (Fig. 5a). Interestingly, the estimated proportion of PT_VCAM1 in the no surgery control mouse kidneys increased in older mice (Fig. 5b) and was accompanied by an increase in leukocytes. These results suggest a role for aging-related chronic inflammation and acute kidney injury in the appearance of PT_VCAM1. To further characterize the role of PT_VCAM1 in acute kidney injury, we used a snRNA-seq mouse IRI^64^ dataset to predict the corresponding cell type for PT_VCAM1 in the injured mouse kidney. Label transfer of cell type annotations from mouse IRI to human indicates that the majority of PT_VCAM1 are related to the failed repair population in the mouse (Failed-repair PT) (Fig. 5c).

**Figure 5.**
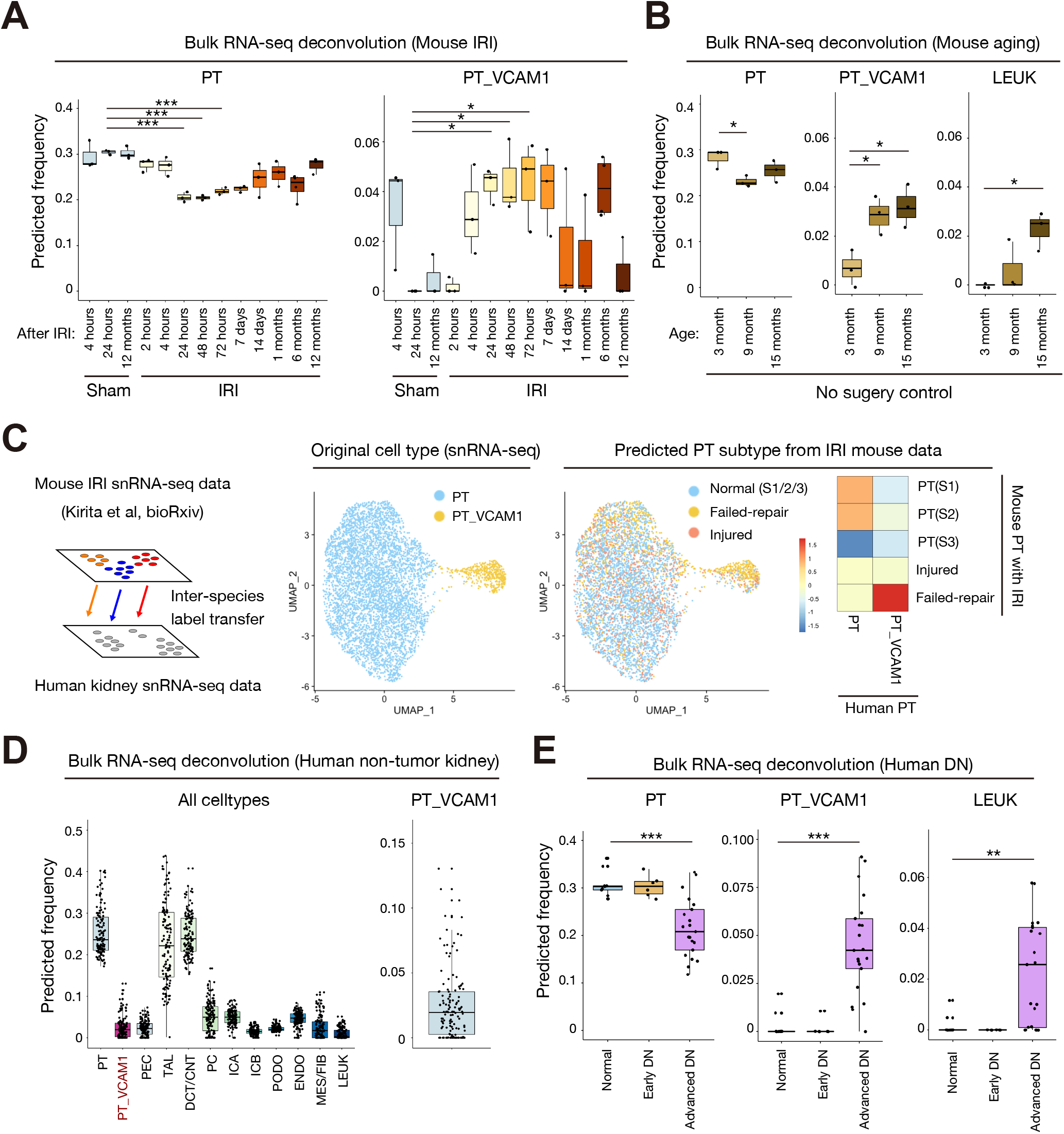
The estimated proportion of VCAM1+ PT increases in acute and chronic kidney disease. (**a**, **b**) Deconvolution analysis of bulk RNA-seq mouse kidney IRI dataset (GSE98622) with BisqueRNA. Sham control and IRI (**a**), or no surgery control (**b**). (**c**) Inter-species data integration was performed between mouse IRI snRNA-seq (GSE139107) and human snRNA-seq with Seurat (left). PT and PT_VCAM1 from human snRNA-seq (middle) are label-transferred from mouse IRI snRNA-seq, and the frequencies of predicted cell types are shown on the heatmap (right). (**d**) Deconvolution analysis of bulk RNA-seq TCGA non-tumor kidney data (**e**) Deconvolution analysis of bulk RNA-seq human diabetic nephropathy (DN) data (GSE142025) with BisqueRNA. Box-and-whisker plots depict the median, quartiles and range. *P < 0.05; **P < 0.01; ***P < 0.005, one-way ANOVA with post hoc Dunnett’s multiple comparisons test.

We retrieved bulk RNA-seq datasets of healthy and injured human kidneys to estimate the proportion of PT_VCAM1^63^. We identified 72 non-tumor kidney samples in The Cancer Genome Atlas (TCGA) using the GDC data portal. The TCGA patients had a mean age of 62.5 years (SD=11.9y) and had undergone nephrectomy for renal cell carcinoma. Deconvolution of the non-tumor kidney samples with BisqueRNA^63^ estimated that the proportion of PT_VCAM1 cells was 2.6% (Fig. 5d), which is consistent with our snRNA-seq and snATAC-seq estimates. Next, we analyzed bulk RNA-seq from kidney biopsies of patients with type 2 diabetes^65^. The patients with advanced diabetic nephropathy had a significantly higher proportion of PT_VCAM1 compared to control or early diabetic nephropathy patients (Fig. 5e), suggesting that PT_VCAM1 may be related to disease progression in diabetic nephropathy.

## Discussion

We performed snRNA-seq and snATAC-seq sequencing in parallel to describe the transcriptional and chromatin accessibility landscape of the adult human kidney. Our analysis demonstrates that snRNA-seq and snATAC-seq are comparable methods for determining cell identity and cell-type-specific chromatin accessibility provides additional information that further elucidates cellular heterogeneity. Multimodal single cell profiling (“multi-omics”)^13^ has greatly improved our ability to detect unique cell types and states while introducing a host of bioinformatics challenges and opportunities^12^. In this study, we outline our integration approach to analyzing paired snRNA-seq and snATAC-seq datasets to highlight functional heterogeneity in the proximal tubule and thick ascending limb.

Studies in mouse and human suggest that there are structural and functional differences between medullary and cortical TAL driven by regional expression patterns of claudins ^47,48^. Claudin-10 and claudin-16 regulate paracellular reabsorption of calcium and magnesium in the thick ascending limb. Mice lacking *Cldn16* develop hypercalciuria and hypomagnesemia, which is similar to the phenotype of patients with familial hypomagnesemia with hypercalciuria and nephrocalcinosis (FHHNC) that carry pathogenic variants in *CLDN16*^44^. In contrast, targeted deletion of *Cldn10* in the thick ascending limb results in impaired paracellular sodium permeability and hypermagnesemia^47^. Interestingly, *Cldn10* deletion can partially rescue the phenotype of *Cldn16-*deficient mice (double knockout)^48^. These data suggest that *Cldn10* and *Cldn16* have opposing effects on cation reabsorption in the thick ascending limb and are supported by the observation that *Cldn10* and *Cldn16* are expressed in a mosaic pattern in mice^66^. We observed two distinct subpopulations of *UMOD+* cells in the thick ascending limb (*CLDN10+CLDN16-* and *CLDN10-CLDN16+*). In motif enrichment analysis, the *CLDN10-CLDN16+* population had increased transcription factor activity of HNF1B (Supplementary Fig.9e,f). Pathogenic germline variants in *HNF1B* are causative of autosomal dominant tubulointerstitial kidney disease with hypomagnesemia and hypercalciuria^51^. Deletion of *Hnf1b* the mouse kidney increases *Cldn10* expression, suggesting that *Cldn10* is regulated by HNF1B transcription factor activity. Future studies using human cell lines derived from TAL or kidney organoids may help to validate this hypothesis.

The proximal tubule is the most abundant cell type in the kidney cortex and is divided into segments (S1, S2, S3) with unique functions driven by segment-specific expression of various transporters, including SGLT1 and SGLT2^67^. SGLT2 is a therapeutic target in diabetic nephropathy and the genes and signaling pathways that regulate SGLT2 expression may be of clinical interest^68^. snRNA-seq detected *SLC5A1* (SGLT1) and *SLC5A2* (SGLT2) in the proximal tubule, but lacked the power to clearly distinguish between the S1/S2 segments that express SGLT2 and the S3 segment that expresses SGLT1. In contrast, snATAC-seq was able to separate the S1/S2 and S3 segments based on chromatin accessibility within the gene body and promoter of *SLC5A1* and *SLC5A2*. These data suggest that snATAC-seq may help to further refine segment-specific cell types; particularly those that are defined by genes transcribed at low levels or genes that are not detected by snRNA-seq. Furthermore, snATAC-seq can predict the transcription factors that drive cell-type-specificity, which may improve our understanding of kidney development and directed differentiation of kidney organoids. We used this approach to implicate NFκB signaling in a subpopulation of proximal tubule epithelial cells.

We used snRNA-seq and snATAC-seq to identify a subpopulation of proximal tubule (PT_VCAM1) that expressed *VCAM1, HAVCR1* (KIM-1), vimentin (*VIM*), *PROM1* (CD133), and *CD24*. The PT_VCAM1 population was also identified in bulk RNA-seq datasets from non-tumor TCGA kidney and human diabetic nephropathy^65^. The proportion of PT_VCAM1 increased in response to acute and chronic kidney injury in both mouse and human^69^. CD133+CD24+ progenitor-like cells have been previously-described in the human kidney in a scattered distribution^18,19^ and VCAM1 (CD106) expression is present in CD133+CD24+ renal progenitors localized to Bowman’s capsule^20^. A separate population of CD133+CD24+CD106-cells are localized to the proximal tubule and both CD133+CD24+CD106+ and CD133+CD24+CD106-cells can engraft in SCID mice to repopulate the tubular epithelium following acute tubular injury^20^. *VCAM1* (*CD106*)*, VIM, PROM1,* and *CD24* expression was enriched in the PT_VCAM1 cluster in our snRNA-seq dataset (Supplementary Fig. 10), which differs from the previously-described CD133+CD24+CD106-renal progenitor population localized to the proximal tubule. We used immunofluorescence studies to demonstrate that VCAM1+ cells are present in a scattered distribution within the proximal tubule of human kidneys (Fig. 4a). Comparison of our human data to a mouse snRNA-seq acute kidney injury dataset ^64^ suggests that PT_VCAM1 is closely-related to a population of ‘failed-repair’ PT, which has a proinflammatory gene expression signature (Fig. 5c). *HAVCR1* expression in PT_VCAM1 suggests that PT_VCAM1 likely represents a subpopulation of PT that is undergoing injury *in situ*, and undergoes expansion in aging and chronic kidney disease (Fig. 5b,d). Pseudotemporal ordering (Fig. 4) indicated that PT_VCAM1 exist along a continuum with PT (Fig. 4), further supporting the hypothesis that they represent an injured cell state. Motif enrichment analysis showed that PT_VCAM1 had increased RELA transcription factor activity and NFκB induction by TNFα increased *VCAM1* expression in an *in vitro* model of proximal tubule cells. Our findings suggest that NFκB plays a role in the maintenance of PT_VCAM1, which may be of clinical interest in designing therapies for acute kidney injury. However, whether proximal tubule repair involves proliferation of a progenitor population or dedifferentiation of mature epithelium still remains controversial^27^ and our own results do not support the existence of a fixed intratubular progenitor population ^70–72^.

An advantage of snATAC-seq is the ability to measure covariance between accessible chromatin sites to predict cis-regulatory interactions^14^. This approach can link putative regulatory regions with their target genes and has been applied to human pancreatic islets^73^, acute leukemia^74^, and multiple mouse tissues, including: hippocampus^75^, mammary gland^76^, T-cells^77^, and kidney among others^8,9^. In particular, genome wide association study (GWAS) risk loci can be linked to their target genes, which would complement the progress made using chromosome conformation capture (Hi-C)^33^. We generated cell-type-specific cis-coaccessibility networks (CCAN) that had significant overlap with a published database^40^. The remaining interactions may represent the unique chromatin interaction landscape of the kidney. We have made all of our data publicly-available and invite readers to explore cell-type-specific differentially accessible chromatin regions (Supplementary Fig. 13).

The small sample size of this study does not adequately capture the expected heterogeneity of the general population. Furthermore, our study focused on kidney cortex and did not include samples from the medulla. Future studies would benefit from studying diseased kidneys to determine how chromatin accessibility changes with progression. Also, improvements in peak calling algorithms for snATAC-seq data will help to narrow the differentially accessible chromatin regions and identify additional peaks in less common cell types. Despite these limitations, our single-cell multimodal atlas of human kidney redefines cellular heterogeneity of the kidney driven by cell-type-specific transcription factors. Our data enhances the understanding of human kidney biology and provides a foundation for future studies.

## Methods

### Tissue procurement

Non-tumor kidney cortex samples were obtained from patients undergoing partial or radical nephrectomy for renal mass at Brigham and Women’s Hospital (Boston, MA) under an established Institutional Review Board protocol. Samples were frozen or retained in OCT for future studies. Histologic sections were reviewed by a renal pathologist and laboratory data was abstracted from the medical record.

### Nuclear dissociation and library preparation

For snATAC-seq, nuclei were isolated with Nuclei EZ Lysis buffer (NUC-101; Sigma-Aldrich) supplemented with protease inhibitor (5892791001; Roche). Samples were cut into < 2 mm pieces, homogenized using a Dounce homogenizer (885302–0002; Kimble Chase) in 2 ml of ice-cold Nuclei EZ Lysis buffer, and incubated on ice for 5 minutes with an additional 2 ml of lysis buffer. The homogenate was filtered through a 40-μm cell strainer (43–50040–51; pluriSelect) and centrifuged at 500g for 5 minutes at 4°C. The pellet was resuspended, washed with 4 ml of buffer, and incubated on ice for 5 minutes. Following centrifugation, the pellet was resuspended in Nuclei Buffer (10× Genomics, PN-2000153), filtered through a 5-μm cell strainer (43–50020–50; pluriSelect), and counted. For snRNA-seq preparation, the RNase inhibitors (Promega, N2615 and Life Technologies, AM2696) were added to the lysis buffer, and the pellet was ultimately resuspended in nuclei suspension buffer (1x PBS, 1% BSA, 0.1% RNase inhibitor) ^78^. 10X Chromium libraries were prepared according to manufacturer protocol.

### Single nucleus RNA sequencing bioinformatics workflow

Five snRNA-seq libraries were obtained using 10X Genomics Chromium Single Cell 5’ v2 chemistry following nuclear dissociation^78^. Three snRNA-seq libraries (patients 1-3) were prepared for a prior study GSE131882^5^. Libraries were sequenced on an Illumina Novaseq instrument and counted with cellranger v3.1.0 using a custom pre-mRNA GTF built on GRCh38 to include intronic reads. Datasets were aggregated with cellranger v3.1.0 without depth normalization and preprocessed with Seurat v3.0.2^12^ to remove low-quality nuclei (Features > 500, Features < 4000, RNA count < 16000, %Mitochondrial genes < 0.8, %Ribosomal protein large or small subunits < 0.4) and DoubletFinder v2.0.2^79^ to remove heterotypic doublets (assuming 5% of barcodes represent doublets). The filtered library was normalized with SCTransform^80^, and corrected for batch effects with Harmony v1.0^81^. After filtering, there was a mean of 3997 +/− 930 cells per snRNA-seq library and a mean of 1674 +/− 913 genes detected per nucleus. Clustering was performed by constructing a KNN graph and applying the Louvain algorithm. Dimensional reduction was performed with UMAP^82^ and individual clusters were annotated based on expression of lineage-specific markers. The final snRNA-seq library contained 19,985 cells and represented all major cell types within the kidney cortex (Supplementary Table 1). Differential expression between cell types was assessed with the Seurat FindMarkers function for transcripts detected in at least 20% of cells.

### Single nucleus ATAC sequencing bioinformatics workflow

Five snATAC-seq libraries were obtained using 10X Genomics Chromium Single Cell ATAC v1 chemistry following nuclear dissociation. Libraries were sequenced on an Illumina Novaseq instrument and counted with cellranger-atac v1.2 (10X Genomics) using GRCh38. Libraries were aggregated with cellranger-atac without depth normalization and processed with Seurat v3.0.2 and its companion package Signac v0.2.1 (https://github.com/timoast/signac)^12^. Low-quality cells were removed from the aggregated snATAC-seq library (peak region fragments > 2500, peak region fragments < 25000, %reads in peaks > 15, blacklist ratio < 0.001, nucleosome signal < 4 & mitochondrial gene ratio < 0.25) before normalization with term-frequency inverse-document-frequency (TFIDF). Dimensional reduction was performed via singular value decomposition (SVD) of the TFIDF matrix and UMAP. A KNN graph was constructed to cluster cells with the Louvain algorithm. Batch effect was corrected with Harmony^81^. A gene activity matrix was constructed by counting ATAC peaks within the gene body and 2kb upstream of the transcriptional start site using protein-coding genes annotated in the Ensembl database^83^. The gene activity matrix was log-normalized prior to label transfer with the aggregated snRNA-seq Seurat object using canonical correlation analysis. The aggregated snATAC-seq object was filtered using a 97% confidence threshold for cell type assignment following label transfer to remove heterotypic doublets. The filtered snATAC-seq object was reprocessed with TFIDF, SVD, and batch effect correction followed by clustering and annotation based on lineage-specific gene activity. After filtering, there was a mean of 5408 +/− 1393 nuclei per snATAC-seq library with a mean of 7538 +/− 2938 peaks detected per nucleus. The final snATAC-seq library contained a total of 214,890 unique peak regions among 27,034 nuclei and represented all major cell types within the kidney cortex (Supplementary Table 2). Differential chromatin accessibility between cell types was assessed with the Signac FindMarkers function for peaks detected in at least 20% of cells using a likelihood ratio test. Genomic regions containing snATAC-seq peaks were annotated with ChIPSeeker^84^ and clusterProfiler^85^ using the UCSC database on hg38^86^.

### Comparison to previously-published database of DNase hypersensitive sites

Glomerulus and tubulointerstitial DNase hypersensitive sites (DHS) were downloaded in bed format from Sieber et al^33^. Glomerulus and tubulointerstitial DHS master lists were composed by merging the tissue-specific bed files, converting to a GRanges object with the GenomicRanges package^87^, and collapsing the intervals with the reduce function. cellranger-atac peaks were filtered by the proportion of nuclei containing the snATAC-seq peak and subsequently overlapped with DHS sites.

### Estimation of transcription factor activity from snATAC-seq data

Transcription factor activity was estimated using the final snATAC-seq library and chromVAR v1.6.0^15^. The positional weight matrix was obtained from the JASPAR2018 database^88^. Cell-type-specific chromVAR activities were calculated using the RunChromVAR wrapper in Signac v0.2.1 and differential activity was computed with the FindMarkers function. Motif enrichment analysis was also performed on the differential accessible regions with the FindMotif function.

### Generation of cis-coaccessibility networks with Cicero

Cis-coaccessibility networks were predicted using the final snATAC-seq library and Cicero v1.2^14^. The snATAC-seq library was partitioned into individual cell types and converted to cell dataset (CDS) objects using the make_atac_cds function. The CDS objects were individually processed using the detect_genes() and estimate_size_factors() functions with default parameters prior to dimensional reduction and conversion to a Cicero CDS object. Cell-type-specific Cicero connections were obtained using the run_cicero function with default parameters.

### Construction of pseudotemporal trajectories with Monocle or Cicero

Monocle3^54^ was used to convert the snRNA-seq dataset into a cell dataset object (CDS), preprocess, correct for batch effects^89^, embed with dimensional reduction and perform pseudotemporal ordering. Cicero ^14^ was used to generate pseudotemporal trajectories for the snATAC-seq dataset. The CDS was constructed from the snATAC-seq dataset, preprocessed, aligned and embedded. The proximal tubular cells identified with Seurat/Signac were designated as the root cells.

### Comparison of Cicero co-accessibility Connections to GeneHancer Database

Cell-type-specific differentially accessible chromatin regions (DAR) were identified with the Signac^12^ FindMarkers function using a log-fold-change threshold of 0.25 for peaks present in at least 20% of cells. Cell-type-specific DAR were extended 50kb up- and downstream to create bed files to query the UCSC table browser^86^ using the GeneHancer interactions tracks^40^. GeneHancer interactions were compared to cell-type-specific Cicero connections to determine the mean proportion of overlap with increasing Cicero coaccess threshold.

### Gene ontology enrichment analysis

Differentially expressed genes in the PT_VCAM1 cluster (compared to PT) were identified with the FindMarkers function using a log-fold-change threshold of 0.25 for the genes expressed in at least 20% of cells. Genes ontology enrichment was performed with PANTHER (http://geneontology.org/)^90,91^.

### Gene set enrichment analysis

Differential expressed genes in PT_VCAM1 cluster (compared to PT) were identified with the FindMarkers function using a log-fold-change threshold of 0.05 for peaks present in at least 5% of cells. The pre-ranked gene list was analyzed with GSEA v4.0.3 (Broad Institute)^60,61^.

### Deconvolution of bulk RNA-seq data

For the TCGA (The Cancer Genome Atlas) dataset, HTseq counts and metadata were downloaded from the GDC data portal (portal.gdc.cancer.gov) by selecting “kidney”, “TCGA”, “RNA-seq”, and “solid tissue normal”. Bulk RNA-seq counts were normalized with DESeq2^92^ and count matrices were deconvoluted with BisqueRNA^63^ using snRNA-seq annotations. For the mouse ischemia reperfusion dataset from Liu et al.^69^, a normalized count matrix was downloaded from GSE98622 and converted to human annotations using biomaRt and ensembl prior to deconvolution with BisqueRNA with default parameters. For the diabetic nephropathy dataset^65^, fastq files were downloaded from GSE128736, transcript abundance was quantified with Salmon using GRCh38, count matrices were imported to DESeq2 with tximport, and data was normalized prior to deconvolution with BisqueRNA.

### Inter-species snRNA-seq data comparison

The snRNA-seq dataset for human adult kidneys was converted to mouse annotations using biomaRt and ensembl, and integrated with a mouse IRI snRNA-seq dataset^64^ using the FindTransferAnchors function in Seurat. Mouse cell type annotations were transferred to the human dataset.

### Renal proximal tubule epithelial cell culture (RPTEC) and TNFα stimulation

RPTEC (Lonza) were cultured with Renal Epithelium Cell Growth Medium 2 (PromoCell). Cells were maintained in a humidified 5% CO_2_ atmosphere at 37°C. Experiments were performed on early passages (passage 2-3). Cells were plated at a density of 1×10^5^ cells per well in a 6-well plate, incubated overnight, and subsequently treated with TNFα (R&D systems, 100 ng/ml). Cells were harvested at 24 h or 48 h after treatment.

### RT and real-time PCR analysis

Total RNA was extracted from RPTEC or kidney organoids with the Direct-zol MicroPrep Kit (Zymo) following manufacturer’s instructions. The extracted RNA (2 ug) was reverse transcribed using the High-Capacity cDNA Reverse Transcription Kit (Life Technologies). Quantitative PCR (RT-PCR) was performed using iTaq Universal SYBR Green Supermix (Bio-Rad). Data were normalized by the abundance of *GAPDH* mRNA. Primer sequences (sense and antisense, respectively) are as follows:

5’- GACAGTCAGCCGCATCTTCT-3’ and 5’- GCGCCCAATACGACCAAATC-3’ for *GAPDH*,
5’- GGGAAGATGGTCGTGATCCTT-3’ and 5’- TCTGGGGTGGTCTCGATTTTA-3’ for *VCAM1*,
5’- GCCGACGTAGCTTCTCTGAAC-3’ and 5’- TTTGGGCTCGACTCTCAATGA-3’ for *TPM1*,
5’- AGGCAACTTCCCGAGAGTTC-3’ and 5’- CCCCAAAGCGGTAGACTTCAG-3’ for *SLC5A12*,
5’- TGATCGGGAGGCTTCTTCTCT-3’ and 5’- GGACCGAAGGTTGGATTTCTTG-3’ for *SLC4A4*.

### Immunofluorescence studies

Formalin-fixed paraffin embedded tissue sections were deparaffinized and underwent antigen retrieval. Sections were blocked with 1% bovine serum albumin, permeabilized with 0.1% Triton-X100 in PBS and incubated overnight with primary antibodies for VCAM1 (abcam, ab134047) and Biotinylated Lotus Tetragonolobus Lectin (Vector Laboratories, B-1325) followed by staining with secondary antibodies (FITC-, Cy3, or Cy5-conjugated, Jackson ImmunoResearch). Sections were stained with DAPI (4′,6′-diamidino-2-phenylindole) and mounted in Prolong Gold (Life Technologies). Images were obtained by confocal microscopy (Nikon C2+ Eclipse; Nikon, Melville, NY).

### Statistical analysis

No statistical methods were used to predetermine sample size. Experiments were not randomized and investigators were not blinded to allocation during library preparation, experiments or analysis. Quantitative data (Fig. 4f) are presented as mean±s.d. and were compared between groups with a two-tailed Student’s *t*-test unless otherwise indicated. Estimated proportion by deconvolution of RNA-seq data (Fig. 5a,b,e) were analysed with one-way ANOVA with post hoc Dunnett’s multiple comparisons test. A *P* value of <0.05 was considered statistically significant.

## Acknowledgments

These experiments were funded by a Chan Zuckerberg Initiative Seed Network Grant and an International Research Fund for Subsidy of Kyushu University School of Medicine Alumni. Y.M. was supported by the Japan Society for the Promotion of Science (JSPS) Postdoctoral Fellowships for Research Abroad, a Postdoctoral Fellowship of Uehara Memorial Foundation and a grant from the Nakatomi Foundation.

## Author contributions

Y.M. and P.C.W. contributed equally to this study. Y.M., P.C.W. and B.D.H. planned the study. S.S.W. collected the kidney samples. Y.M. prepared the snRNA-seq and snATAC-seq libraries. Y.M., P.C.W. and B.D.H. analysed and interpreted snRNA-seq and snATAC-seq data. Y.M. performed other experiments. H.W. generated the public database for data. Y.M., P.C.W. and B.D.H. wrote the manuscript. B.D.H. coordinated and oversaw the study. All authors discussed the results and commented on the manuscript.

