## Supplemental Materials for "Single cell transcriptional and chromatin accessibility profiling redefine cellular heterogeneity in the adult human kidney"

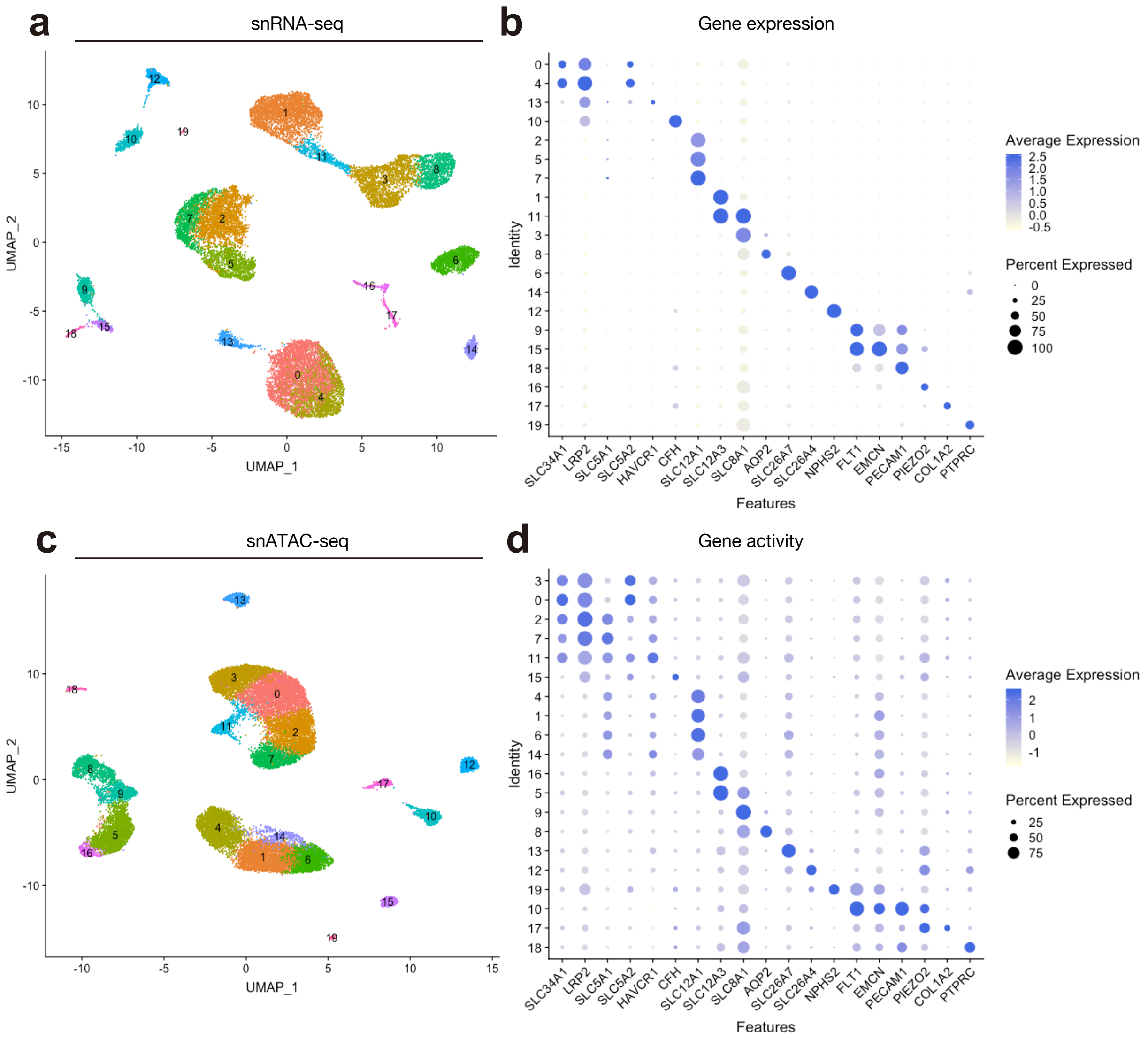

**Supplementary Figure 1 – Unsupervised clustering of snRNA-seq and snATAC-seq human kidney datasets:** (**a**) Unsupervised clustering of snRNA-seq dataset in Seurat, and (**b**) the dot plots showing marker gene expressions of each cell types. (**c**) Unsupervised clustering of snATAC-seq dataset in Signac, and (**d**) the dot plots showing marker gene activities of each cell types.

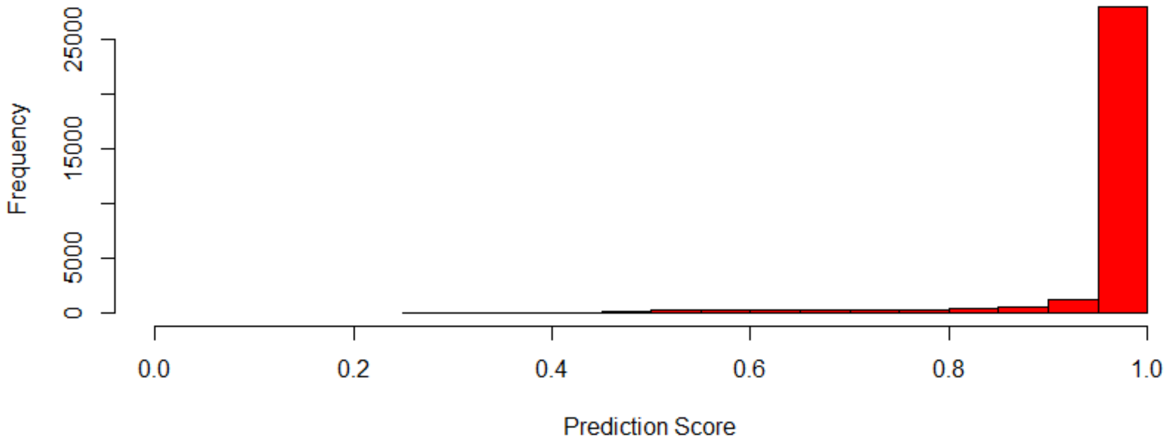

**Supplementary Figure 2 – Label transfer of annotated snRNA-seq confidently predicts snATAC-seq cell types:** Distribution of maximum prediction scores of nuclei calculated by the label transfer algorithm in Signac package. A gene activity matrix was created from the snATAC-seq data and transfer anchors were identified between the 'reference' snRNA-seq dataset and 'query' gene activity matrix followed by assignment of predicted cell types using the Signac package.

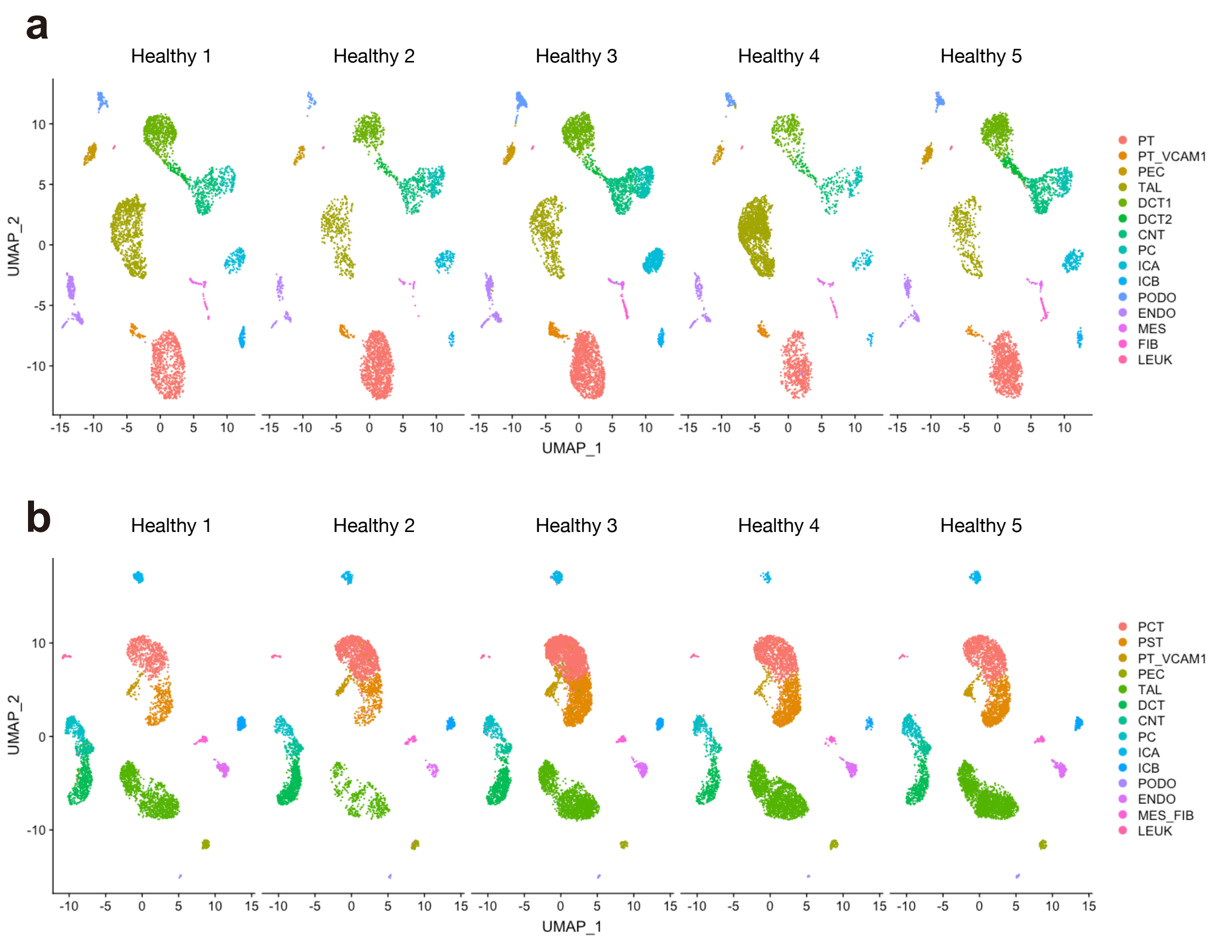

**Supplementary Figure 3 – All cell types detected are found in each kidney sample in both modalities:** (**a**) UMAP visualization of snRNA-seq dataset per kidney sample. (**b**) UMAP visualization of snATAC-seq dataset per kidney sample.

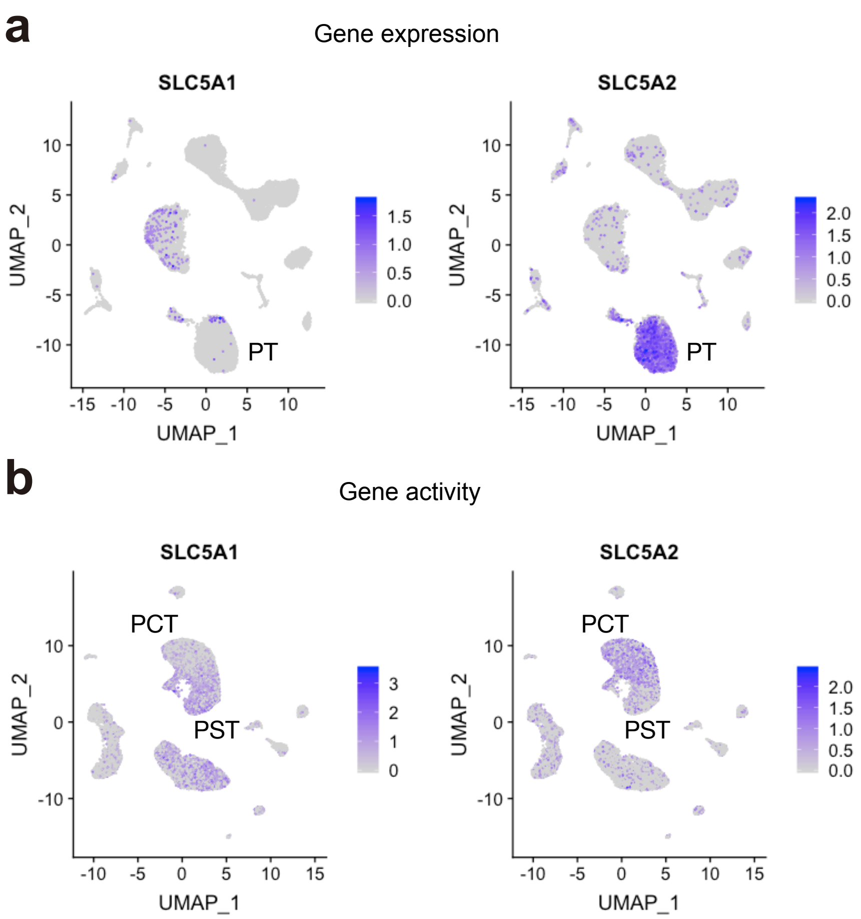

**Supplementary Figure 4 – SLC5A1 and SLC5A2 gene activity delineate proximal tubule segments:** (**a**) snRNA-seq does not detect *SLC5A1* expression, and *SLC5A2* expression does not clearly distinguish subpopulations of proximal tubule. (**b**) snATAC-seq shows increased *SLC5A1* gene activity in a subpopulation of proximal tubule (PST) that is mutually exclusive to the subpopulation (PCT) showing increased *SLC5A2* gene activity.

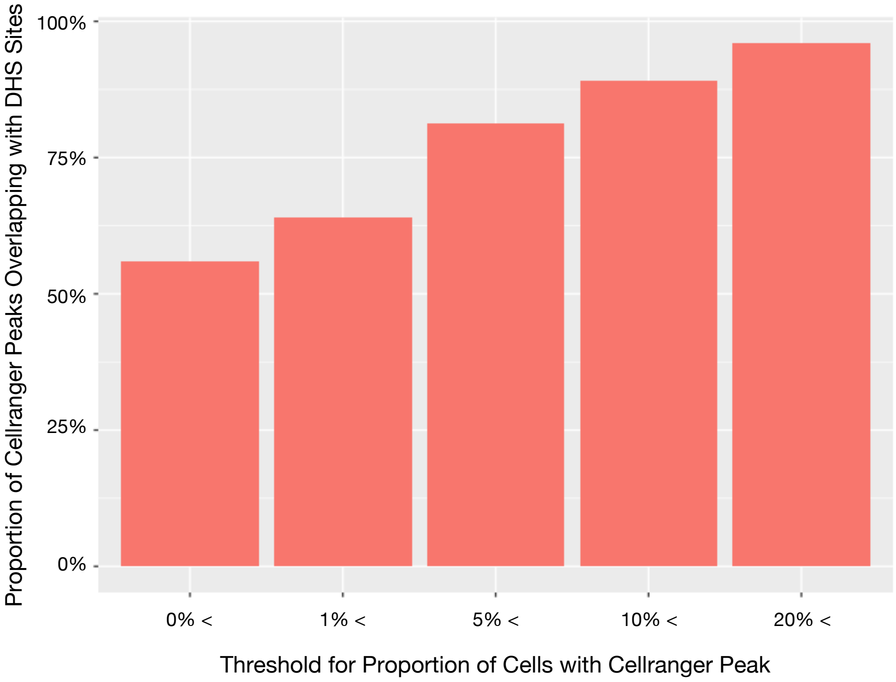

**Supplementary Figure 5 – Aggregated Cell Ranger peaks significantly overlap with previously-published DNase hypersensitive sites:** Cell Ranger peaks were filtered for peaks contained in a designated proportion of cells (x-axis). DNase hypersensitive sites (DHS) were downloaded from Sieber et al. (PMID: 30760496) and overlapped with the Cell Ranger peaks using the GenomicRanges package. The proportion of overlap between DHS and Cell Ranger peaks increased as Cell Ranger peaks were filtered for peaks present in an increasing proportion of cells.

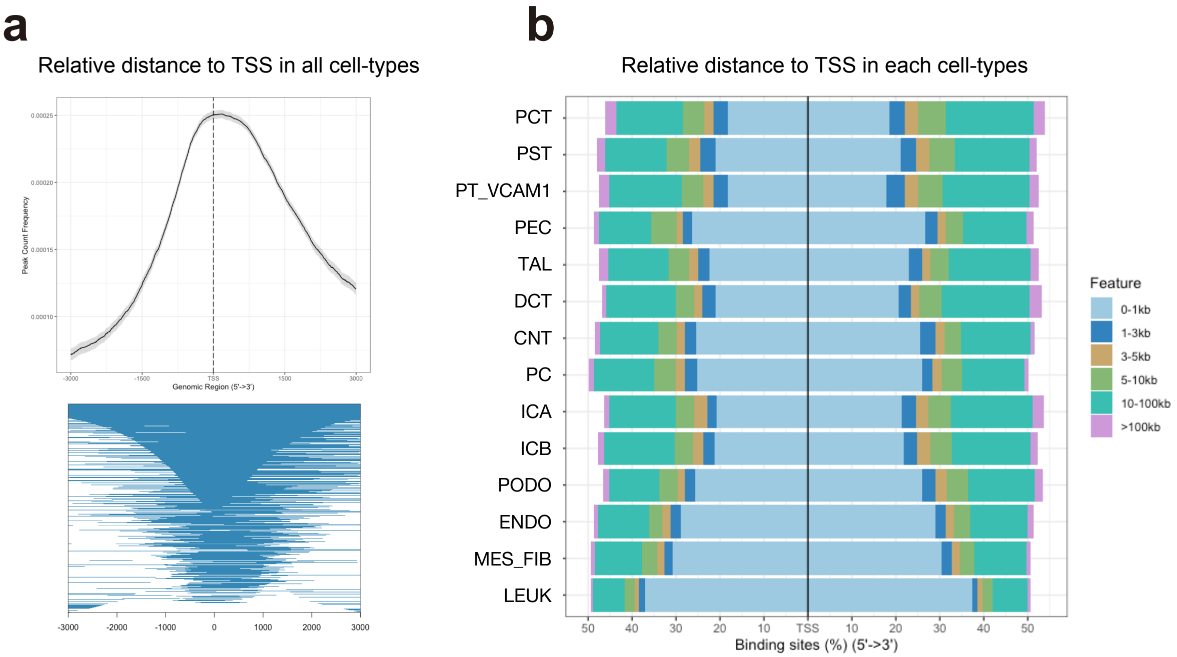

**Supplementary Figure 6 – Cell type-specific DAR are enriched around the transcription start sites (TSS):** (**a**) Relative distance of differentially accessible region (DAR) to TSS in all the dataset. (**b**) Relative distance of DAR to TSS in each cell type.

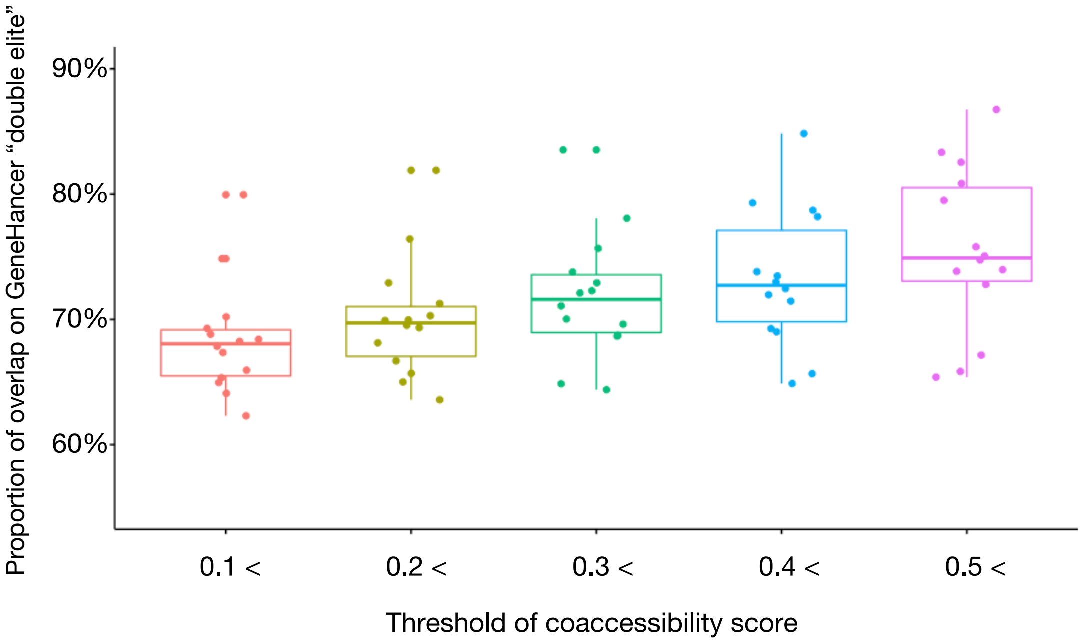

**Supplementary Figure 7– Cicero connections significantly overlap with the GeneHancer interaction database:** The snATAC-seq dataset was partitioned into individual cell types and cell-type-specific cis-coaccessibility networks (CCAN) were identified with the R package Cicero. Cicero connections within 50kb of a cell-type-specific differentially accessible region (DAR) were compared to GeneHancer 'double elite' interactions downloaded from the UCSC table browser for varying Cicero coaccessibility thresholds, and the percentage of overlapped interactions are shown. Each dot is each cell type. Box-and-whisker plots depict the median, quartiles and range.

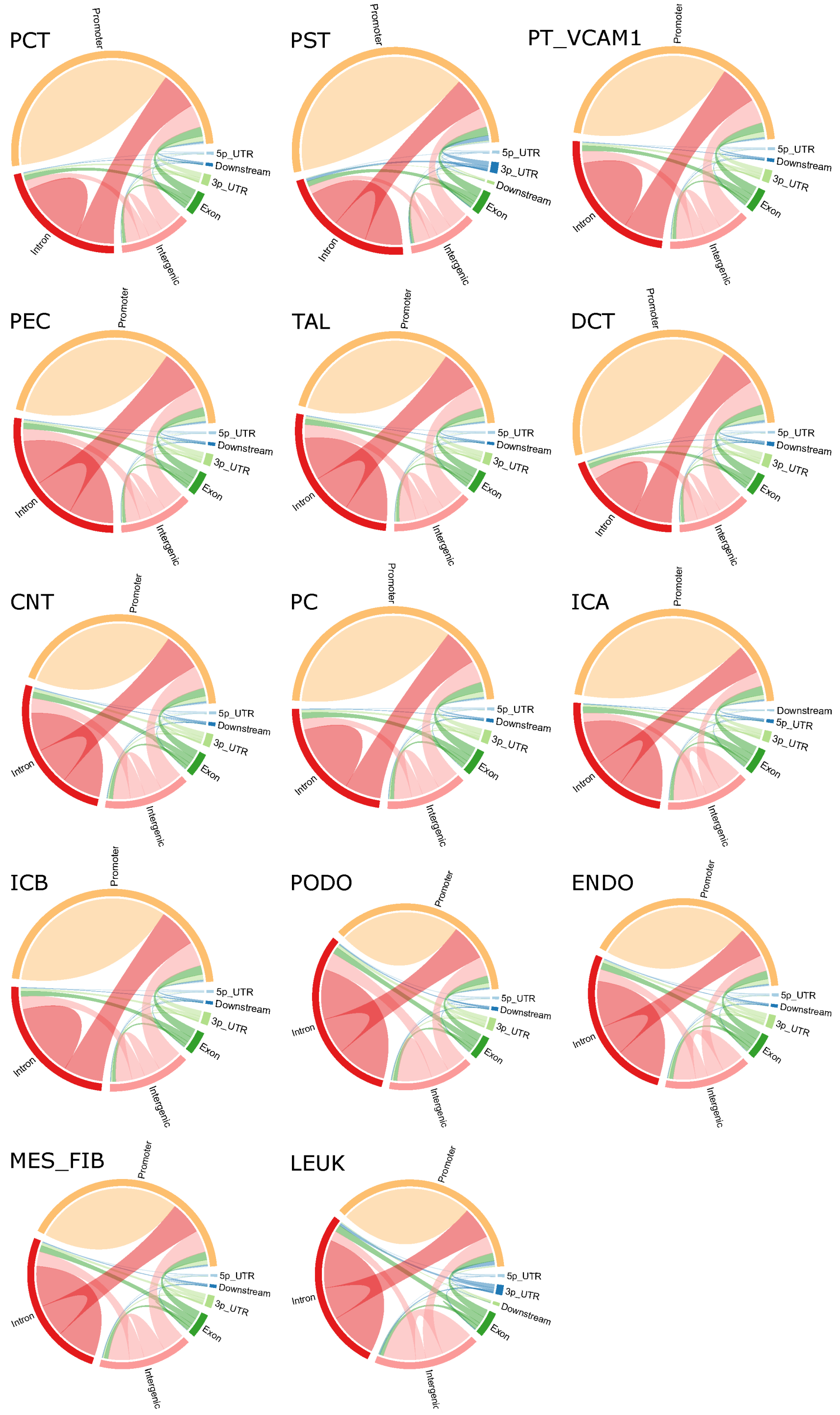

**Supplementary Figure 8 – Annotation of Cicero Connections:** The snATAC-seq dataset was partitioned into individual cell types and cis-coaccessibility networks were predicted with Cicero. The Cicero connection endpoints with a coaccessibility threshold > 0.2 were annotated with ChIPSeeker using the UCSC database. The relative number of connections within and between the designated genomic regions is displayed for each cell type.  Promoter - region within 3kb of the transcriptional start site. 3p_UTR- 3’ untranslated region, 5p_UTR- 5’ untranslated region, Downstream - 3kb downstream of the 3’ UTR.

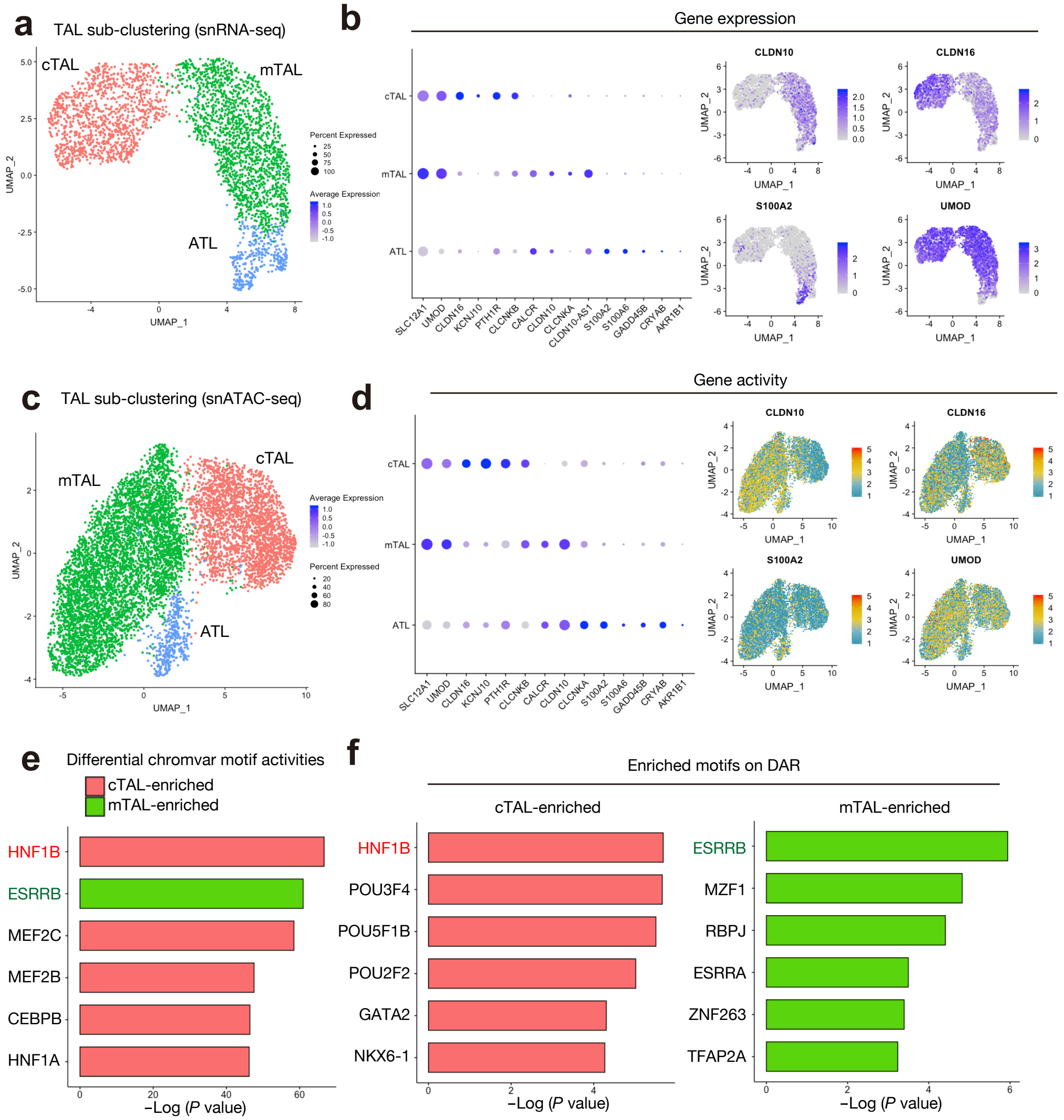

**Supplementary Figure 9 – Transcriptional and epigenetic heterogeneity in the thick ascending limb:**

(**a**) Sub-clustering of TAL on the umap plot of snRNA-seq dataset to divide three subpopulations (cTAL, mTAL and ATL). cTAL, cortical TAL; mTAL, medullary TAL; ATL, Ascending thin limb (of loop of Henle). (**b**) Dotplots showing gene expression patterns of the genes enriched in each of TAL subpopulations (left). Umap displaying gene expressions of *CLDN10*, *CLDN16*, *S100A2* or *UMOD* (right). (**c**) Sub-clustering of TAL on the umap plot of snATAC-seq dataset to divide three subpopulations (cTAL, mTAL and ATL). (**d**) Dotplots showing gene activity patterns of the genes enriched in each of TAL subpopulations (right). Umap displaying gene activities of *CLDN10*, *CLDN16*, *S100A2* or *UMOD* (left). (**e**) Differentially activated transcription factor motifs with chromVAR between cTAL and mTAL. The top 6 motifs with the lowest p values are listed. (**f**) Motif enrichment analysis on the DARs between cTAL and mTAL. Background was set as the genomic regions that are accessible to at least 2.5% of the TAL cells. The top 6 motifs with the lowest p values in each subpopulation are listed.

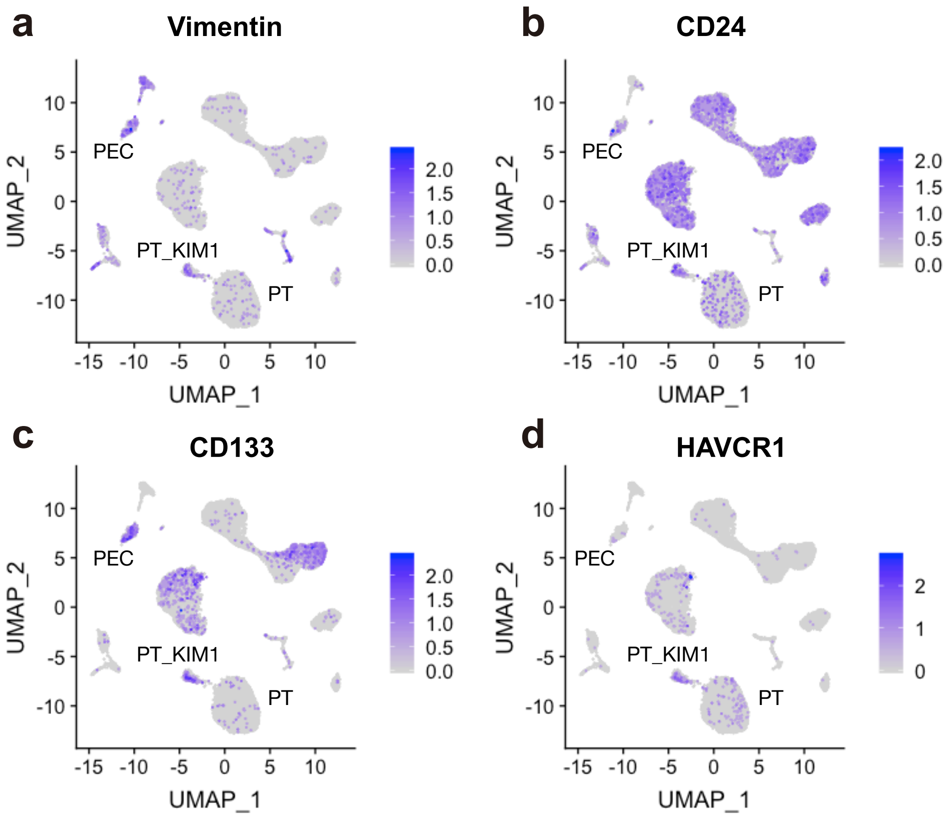

**Supplementary Figure 10 – Marker gene expressions in the PT_VCAM1 population**: (**a**) Vimentin (**b**) CD24, (**c**) CD133 and (**d**) HAVCR1 expression in the snRNA-seq dataset shows increased expression of these genes in the PT_VCAM1 population compared to PT.

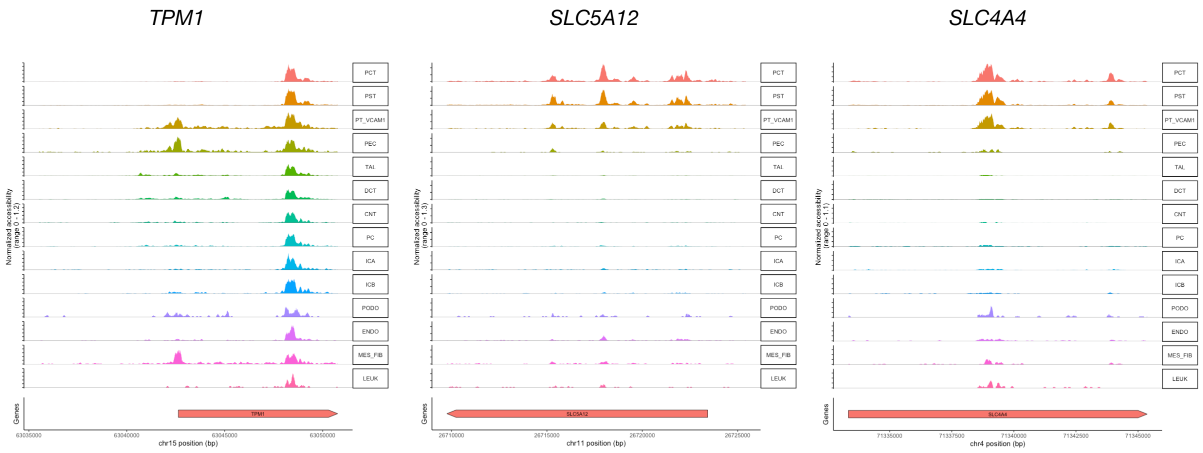

**Supplementary Figure 11 – Fragment coverage around representative DAR in PT_VCAM1:** Fragment coverage (frequency of Tn5 insertion) around the representative DAR (DAR +/-5000 bp) on *TPM1*, *SLC5A12* or *SLC4A4* locus shown in Fig.4D.

**a**

| Hallmark gene set enriched in PT_VCAM1 | NES | *P*-val | FDR-*q* |
|---|---|---|---|
| TNFA_SIGNALING_VIA_NFKB | 2.01 | 0.000 | 0.011 |
| INTERFERON_GAMMA_RESPONSE | 1.84 | 0.002 | 0.070 |
| HYPOXIA | 1.79 | 0.003 | 0.112 |
| UV_RESPONSE_DN | 1.77 | 0.002 | 0.127 |
| EPITHELIAL_MESENCHYMAL_TRANSITION | 1.76 | 0.003 | 0.148 |

**b**

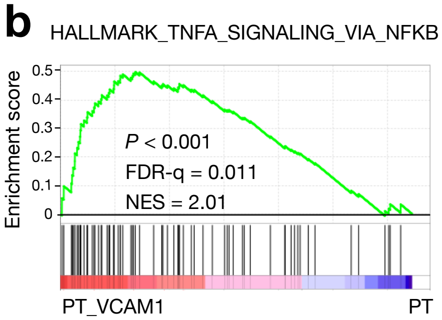

**Supplementary Figure 12 – Gene set enrichment analysis on the differential expressed genes in PT_VCAM1 vs PT suggested activation of NFKB pathway genes**: GSEA of differentially expressed genes for hallmark gene sets (**a**) and the HALLMARK_TNFA_SIGNALING_VIA_NFKB gene set (genes regulated by NFκB induced by TNFα) (**b**) in PT_VCAM1 compared with PT.

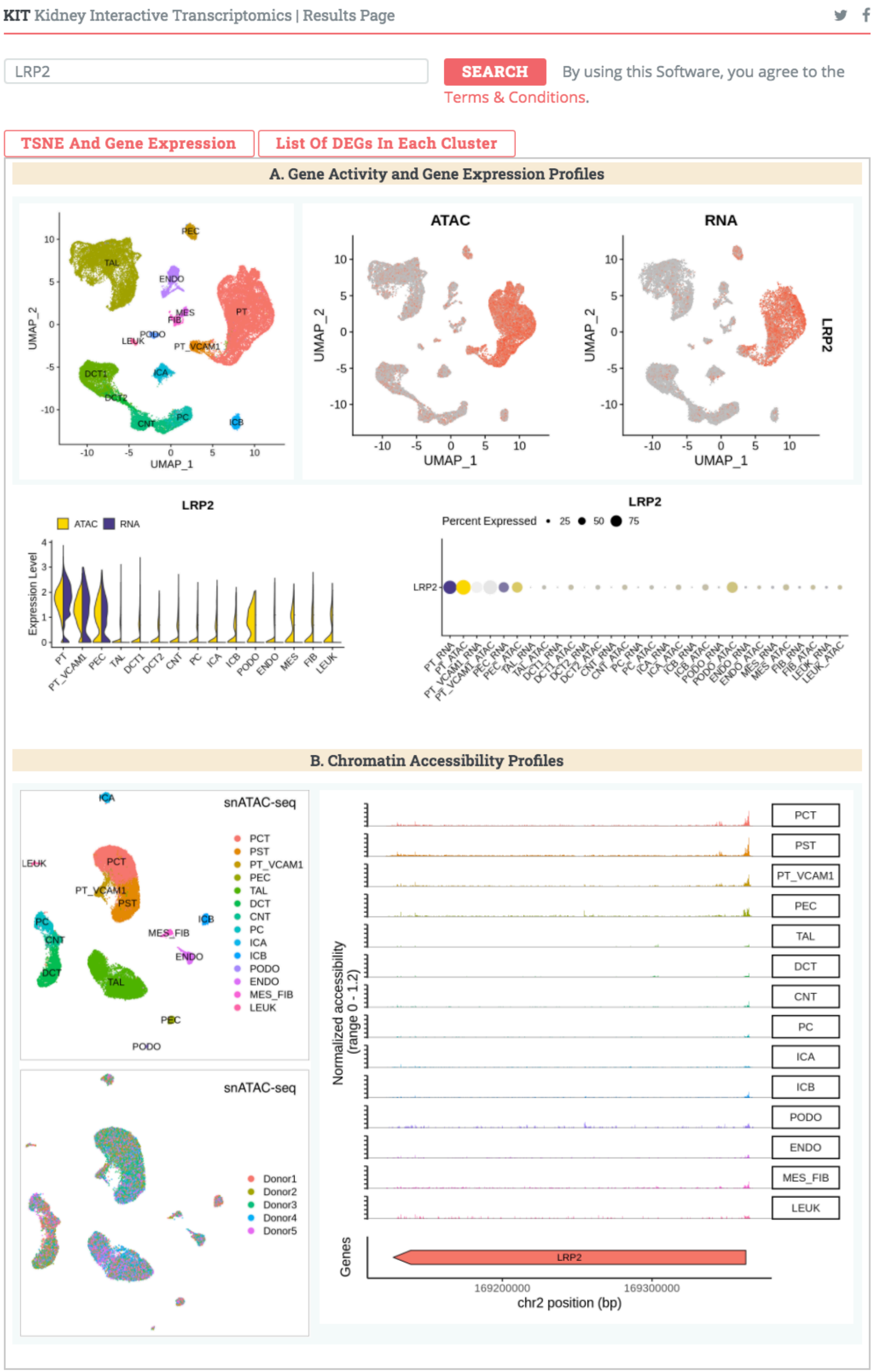

**Supplementary Figure 13 – Online analyzer for the harmonized multimodal kidney cell atlas encompassing both transcriptomic and epigenomic data**: Cell type-specific differential expressed genes, chromatin accessibilities, gene activities and predicted transcription factor motif activities are searchable on the webpage (http://humphreyslab.com/SingleCell/).

| Supplementary Table 1 – Patient Demographics, Laboratory Data, and Renal Pathology | | | | | | | | |
|---|---|---|---|---|---|---|---|---|
| ID | Age | Race | Sex | eGFR (ml/min/1.73m$^2$) | sCr (mg/dL) | Glomerulosclerosis | IFTA | ANS |
| Healthy 1 | 54 | NHW | M | 58 | 1.28 | None, < 10% | 1-10% | Mild |
| Healthy 2 | 62 | HIS | M | 61 | 1.21 | None, < 10% | 1-10% | Moderate |
| Healthy 3 | 61 | NHW | F | 69 | 0.89 | None, < 10% | 1-10% | Mild |
| Healthy 4 | 50 | NHW | M | 78 | 1.10 | None, < 10% | 1-10% | Moderate |
| Healthy 5 | 52 | NHW | F | 98 | 0.89 | None, < 10% | 1-10% | Mild |

**Supplementary Table 1 – Patient demographics and clinical information abstracted from the medical record:** Histologic review was performed by a renal pathologist. NHW-non-hispanic white, HIS-hispanic or latino, IFTA-interstitial fibrosis and tubular atrophy, ANS-arterial and arteriolar nephrosclerosis.

| **Supplementary Table 2**<br>**Filtered snRNA-seq Dataset**<br>**Cell Identity** | **Number** | **Frequency** |
|---|---|---|
| PT | 5036 | 25.2% |
| PT_VCAM1 | 449 | 2.2% |
| PEC | 552 | 2.8% |
| TAL | 4435 | 22.2% |
| DCT1 | 2761 | 13.8% |
| DCT2 | 489 | 2.4% |
| CNT | 1805 | 9.0% |
| PC | 1022 | 5.1% |
| ICA | 1107 | 5.5% |
| ICB | 349 | 1.7% |
| PODO | 463 | 2.3% |
| ENDO | 1008 | 5.0% |
| MES | 239 | 1.2% |
| FIB | 207 | 1.0% |
| LEUK | 63 | 0.3% |
| **TOTAL** | **19985** | 100% |

**Supplementary Table 2 – The number or frequency of cells for each cell type quantitated in the filtered snRNA-seq dataset:** PT-proximal tubule, PT_VCAM1-proximal tubule VCAM1+, PEC-parietal epithelial cells, TAL-thick ascending limb, DCT1-distal convoluted tubule segment 1, DCT2-distal convoluted tubule segment 2, CNT-connecting tubule, PC-principal cells, ICA-intercalated cells type A, ICB-intercalated cells type B, PODO-podocytes, ENDO-endothelial cells, MES-mesangial cells, FIB-fibroblasts, LEUK-leukocytes

| Supplementary Table 3<br>Filtered snATAC-seq Dataset | | |
|---|---|---|
| **Cell Identity** | **Number** | **Frequency** |
| PCT | 6268 | 23.2% |
| PST | 4280 | 15.8% |
| PT_VCAM1 | 674 | 2.5% |
| PEC | 403 | 1.5% |
| TAL | 7762 | 28.7% |
| DCT | 2777 | 10.3% |
| CNT | 898 | 3.3% |
| PC | 1302 | 4.8% |
| ICA | 611 | 2.3% |
| ICB | 620 | 2.3% |
| PODO | 135 | 0.5% |
| ENDO | 759 | 2.8% |
| MES_FIB | 352 | 1.3% |
| LEUK | 193 | 0.7% |
| **TOTAL** | **27034** | 100% |

**Supplementary Table 3 – The number of cells or frequency for each cell type quantitated in the filtered snATAC-seq dataset:** PCT-proximal convoluted tubule, PST-proximal straight tubule, PT_VCAM1-proximal tubule VCAM1+, PEC-parietal epithelial cells, TAL-thick ascending limb, DCT-distal convoluted tubule, CNT-connecting tubule, PC-principal cells, ICA-intercalated cells type A, ICB-intercalated cells type B, PODO-podocytes, ENDO-endothelial cells, MES_FIB-mesangial cells and fibroblasts, LEUK-leukocytes.

| **Supplementary Table 4: Overlap between Cell-type-specific differentially expressed genes and accessible chromatin regions** | | | | | | |
|---|---|---|---|---|---|---|
| **snRNA_vs_snATAC** | **# DEG** | **DEG with DAR** | **Prop. DEG with DAR** | **# DAR** | **DAR near DEG** | **Prop. DAR near DEG** |
| PT_vs_PCT | 769 | 333 | 0.43 | 3055 | 618 | 0.20 |
| PT_vs_PST | 769 | 263 | 0.34 | 2273 | 439 | 0.19 |
| PT_VCAM1 | 425 | 128 | 0.30 | 1315 | 201 | 0.15 |
| PEC | 627 | 190 | 0.30 | 1221 | 267 | 0.22 |
| TAL | 408 | 169 | 0.41 | 1704 | 262 | 0.15 |
| DCT1_vs_DCT | 432 | 178 | 0.41 | 1401 | 277 | 0.20 |
| DCT2_vs_DCT | 348 | 142 | 0.41 | 1401 | 230 | 0.16 |
| CNT | 442 | 123 | 0.28 | 1146 | 185 | 0.16 |
| PC | 523 | 169 | 0.32 | 1416 | 268 | 0.19 |
| ICA | 651 | 236 | 0.36 | 1427 | 379 | 0.27 |
| ICB | 648 | 251 | 0.39 | 1754 | 421 | 0.24 |
| PODO | 927 | 335 | 0.36 | 1712 | 526 | 0.31 |
| ENDO | 861 | 421 | 0.49 | 2781 | 699 | 0.25 |
| MES_vs_MES_FIB | 774 | 167 | 0.22 | 1203 | 231 | 0.19 |
| FIB_vs_MES_FIB | 741 | 179 | 0.24 | 1203 | 248 | 0.21 |
| LEUK | 846 | 396 | 0.47 | 3642 | 611 | 0.17 |
| min | 348 | 123 | 0.22 | 1146 | 185 | 0.15 |
| max | 927 | 421 | 0.49 | 3642 | 699 | 0.31 |
| mean | 636.94 | 230.00 | 0.36 | 1790.88 | 366.38 | 0.20 |
| stdev | 185.61 | 94.92 | 0.08 | 752.23 | 166.80 | 0.04 |

**Supplementary Table 4 – Overlap between cell-type-specific differentially expressed genes and accessible chromatin regions:** Cell-type-specific differentially expressed genes (DEG) were identified for each cell type in the snRNA-seq dataset using the Seurat FindAllMarkers function with a log-fold threshold of 0.25 for genes expressed in at least 20% of cells. Cell-type-specific differentially accessible chromatin regions (DAR) were identified for each cell type in the snATAC-seq dataset using the Signac FindAllMarkers function with a log-fold threshold of 0.25 for peaks present in at least 20% of cells. DAR were annotated with the closest gene in the Ensembl database. The annotated gene list was overlapped with DEG to determine the proportion of DEG with a nearby DAR (Prop. DEG with DAR) and the proportion of DAR with a nearby DEG (Prop. DAR near DEG).
